# Structure of the RPAP3:TRBP interaction reveals an involvement of the HSP90/R2TP chaperone complex in dsRNA pathways

**DOI:** 10.1101/2020.11.11.367672

**Authors:** Yoann Abel, Christophe Charron, Valérie Bourguignon-Igel, Marc Quinternet, Marie-Eve Chagot, Céline Verheggen, Christiane Branlant, Edouard Bertrand, Xavier Manival, Bruno Charpentier, Mathieu Rederstorff

**Author notes:** IGMM, CNRS, Université de Montpellier, Montpellier, 34293, France. These authors should be considered as co-authors (Yoann Abel, Christophe Charron).

## Abstract

MicroRNAs silence mRNAs by guiding the RISC complex. RISC assembly requires cleavage of pre-miRNAs by Dicer, assisted by TRBP or PACT, and the transfer of miRNAs to AGO proteins. The R2TP complex is an HSP90 cochaperone involved in the assembly of ribonucleoprotein particles. Here, we show that the R2TP component RPAP3 binds TRBP but not PACT. Specifically, the RPAP3-TPR1 domain interacts with the TRBP-dsRBD3 and the 1.5 Å resolution crystal structure of this complex is presented. We identify key residues involved in the interaction and show that binding of TRBP to RPAP3 or Dicer is mutually exclusive. In contrast, RPAP3 can simultaneously bind TRBP and HSP90. Interestingly, AGOs and Dicer are sensitive to HSP90 inhibition and TRBP becomes sensitive in absence of RPAP3. These data indicate that the HSP90/R2TP chaperone is an important cofactor of proteins involved in dsRNA pathways.

## INTRODUCTION

MicroRNAs (miRNAs) play essential roles in regulating gene expression. Their biogenesis begins in the nucleus with the processing of a pri-miRNA by the microprocessor complex, composed of the type III Ribonuclease (RNase) Drosha and its cofactor DGCR8 (DiGeorge syndrome Critical Region 8), giving rise to the pre-miRNA. After cytoplasmic export via the exportin 5/Ran-GTP pathway, the pre-miRNA is further processed into the mature, double-stranded miRNA duplex by the cytoplasmic RNase III Dicer (1,2), which is associated to one of its two double-stranded RNA binding protein co-factors, TRBP (TransActivation Response –TAR– RNA binding protein) or PACT (protein activator of the double-stranded RNA-dependent kinase-PKR). Finally, one strand of the cleaved pre-miRNA is loaded onto an Argonaute protein (Ago) in the RNA induced silencing complex (RISC) (2).

As illustrated by their names, prior to their functions as co-factors of Dicer, both TRBP and PACT proteins were initially identified for their positive and negative roles in HIV infection, respectively. Indeed, TRBP has several positive effects on HIV multiplication. It was initially identified as a binding factor of the TAR RNA element of human immunodeficiency viruses HIV-1 and 2 (3). The 5’-terminal TAR stem-loop structure of HIV RNAs impedes efficient translation of the viral RNAs (4) and TRBP binding to this element relieves this negative effect. Interestingly, Dicer was also recently proposed to be involved in this process (5). TRBP was additionally shown to promote HIV infection by directly or indirectly inhibiting PKR activation, which is triggered by the TAR RNA and leads to global translation inhibition (6). More precisely, TRBP inhibits PKR activity via a direct interaction that is reinforced when TRBP is phosphorylated (7,8). Furthermore, PKR activity can also be impeded by TRBP through its binding to PACT, which prevents the latter’s activating interaction with PKR. Finally, another important activity of TRBP in favor of HIV multiplication was recently discovered: TRBP recruits the 2’-*O*-methyltransferase FTSJ3 on HIV RNA (9), which subsequently methylates the viral genome at several specific positions enabling viral escape from the host’s innate immune response.

TRBP and PACT are both composed of three double-stranded RNA binding domains (dsRBD), the two first ones are involved in double-stranded RNA (dsRNA) binding and classified as canonical type A dsRBDs, while the third one mediates protein-protein interactions, in particular with Dicer, and corresponds to a non-canonical type B dsRBD (10,11). While TRBP contributes both to pre-miRNAs and pre-siRNAs processing by Dicer, PACT participates more efficiently to pre-miRNA processing, a specificity that was shown to be mediated by the N-terminal domain of the two cofactors (12). Additionally, association of Dicer to TRBP or PACT was shown to generate miRNAs of different sizes, and possibly of different target repertoire, referred to as isomiRs (10,13). Once the pre-miRNA has been cleaved in the cytoplasm, one of the single-strand derived from the mature miRNA duplex is loaded onto the AGO2 protein within the RNA induced silencing complex (RISC), with the help from the HSC70/HSP90 chaperone machinery (14). This machinery was shown to stabilize free AGO2 and to stimulate efficient miRNA loading, but was also shown not to be involved in target cleavage or inhibition of translation (14–16). Interestingly, the HSC70/HSP90 chaperones have numerous co-chaperones (17). Of particular interest is the R2TP co-chaperone complex, playing a crucial role in the assembly and maturation of large macromolecular complexes essential for most of the universally conserved nanomachines of eukaryotic cells (18–20). This includes several RNPs, such as the U4 and U5 snRNPs, telomerase, as well as the C/D and H/ACA snoRNPs involved in ribosome biogenesis (18,19,21–25). It also includes protein-only clients, such as the nuclear RNA polymerase II (26,27), dynein (28) or complexes containing any of the phosphatidylinositol 3-kinase–like family of kinases (PIKKs): mammalian Target Of Rapamycin (mTOR; (29,30), ATM and RAD3-related (ATR) interacting protein (ATRIP) (31), Suppressor with Morphogenetic effect on G*enitalia* (SMG1; (32), DNA-PK and TRRAP (33).

The R2TP complex consists of a RPAP3:PIH1D1 heterodimer associated to a hetero-hexamer of RUVBL1 and RUVBL2, which are related AAA+ ATPases that also display chaperone activities (22). In metazoans, the R2TP is part of a larger chaperone complex called the PAQosome, which contains an additional series of prefoldin-like proteins and POLR2E and WDR92 (34). Within R2TP, PIH1D1 is believed to play important roles in specifying and recruiting clients, in part via its ability to specifically bind CK2 phosphorylation sites, i.e. phosphoserines embedded in acidic regions of DSDD/E consensus (29,35). RPAP3 regulates HSP90 activity (36,37) and also plays a scaffolding role as it makes stable interactions with all the other components of the R2TP complex (36). It binds HSP90 with its two TPR domains, PIH1D1 via a small peptide sequence located immediately after the TPRs, and the RUVBL1/2 heterohexamer with its conserved C-terminal domain (38–41). However, RPAP3 has not so far been involved in client recognition.

Here, we identified a direct interaction between the TPR1 domain of RPAP3 and the dsRBD3 of TRPB (10,42), and showed that this interaction is exclusive from that between TRBP and Dicer. The X-ray structure of the TRBP/RPAP3 complex at a resolution of 1.5 Å brings exciting novel insights towards understanding structural and molecular aspects of chaperones in the dsRNA pathways. HSP90 inhibition tests revealed putatively novel functions for the R2TP complex in stabilizing dsRNA-pathways proteins.

## RESULTS

### Human TRBP interacts with RPAP3 *in vitro* and in human cells

To determine whether the R2TP complex might be linked to miRNP or RISC assembly, and to identify putatively novel protein/protein interactions between co-chaperones and components of the miRNA biogenesis machinery, we performed a candidate-based yeast two-hybrid (Y2H) screen in *S. cerevisiae*. Interestingly, we found that RPAP3, a member of the R2TP complex, efficiently associated with TRBP, one of the two cofactors of Dicer **(Fig. 1 a, left and middle panels, Supp. Fig. 1 a, b)** (12), but not with PACT, despite a similar overall structural organization **(Fig. 1 a, right panel, Supp. Fig. 1 b-g)**. The association between RPAP3 and TRBP appeared rather strong, as diploid cells grew at a concentration of 3-AT up to 40 mM, which is comparable to the positive control association between Dicer and TRBP **(Fig. 1 a, left and middle panels)**. As interactions detected by Y2H assays might be mediated by additional factors, we performed co-expression experiments in *E. coli* to test whether the TRBP:RPAP3 interaction was direct **(Fig. 1 b, Supp. Fig. 2 a, b)**. At low salt conditions (50 mM NaCl), RPAP3 copurified with a His_6_-tagged version of TRBP using cobalt beads (TALON; **Fig. 1 b, lane 5)**. Additionally, using *in vitro* assays with purified recombinant proteins and purification on glutathione beads, the His_6_-tagged TRBP protein copurified with GST (glutathione-S-transferase)-tagged RPAP3, but not with GST alone **(Supp. Fig. 3 a)**. Because we validated these interactions using recombinant proteins expressed in *E. coli*, we additionally validated the interaction between RPAP3 and TRBP by coimmunoprecipitation in human embryonic kidney cells. We observed that RPAP3 coimmunoprecipitated with endogenously expressed flagged TRBP protein in a doxycycline inducible HEK293 T-Rex cell line **(Fig. 1 c)**, as well as with a transiently expressed V5 tagged-TRBP, even in cell lysates treated with RNase A, suggesting that the interaction was not mediated by RNA **(Supp. Fig. 3 b)**. Finally, we validated the interaction of both endogenous TRBP and RPAP3 *in cellulo* using a proximity ligation assay in HeLa cells **(Duolink; Fig. 1 d, RPAP3:TRBP: PLA and merge; Supp. Fig. 3 c for controls)**. Based on the overall data, we concluded that RPAP3 directly interacts with TRBP, both *in vitro* and *in cellulo*.

**Figure 1.**
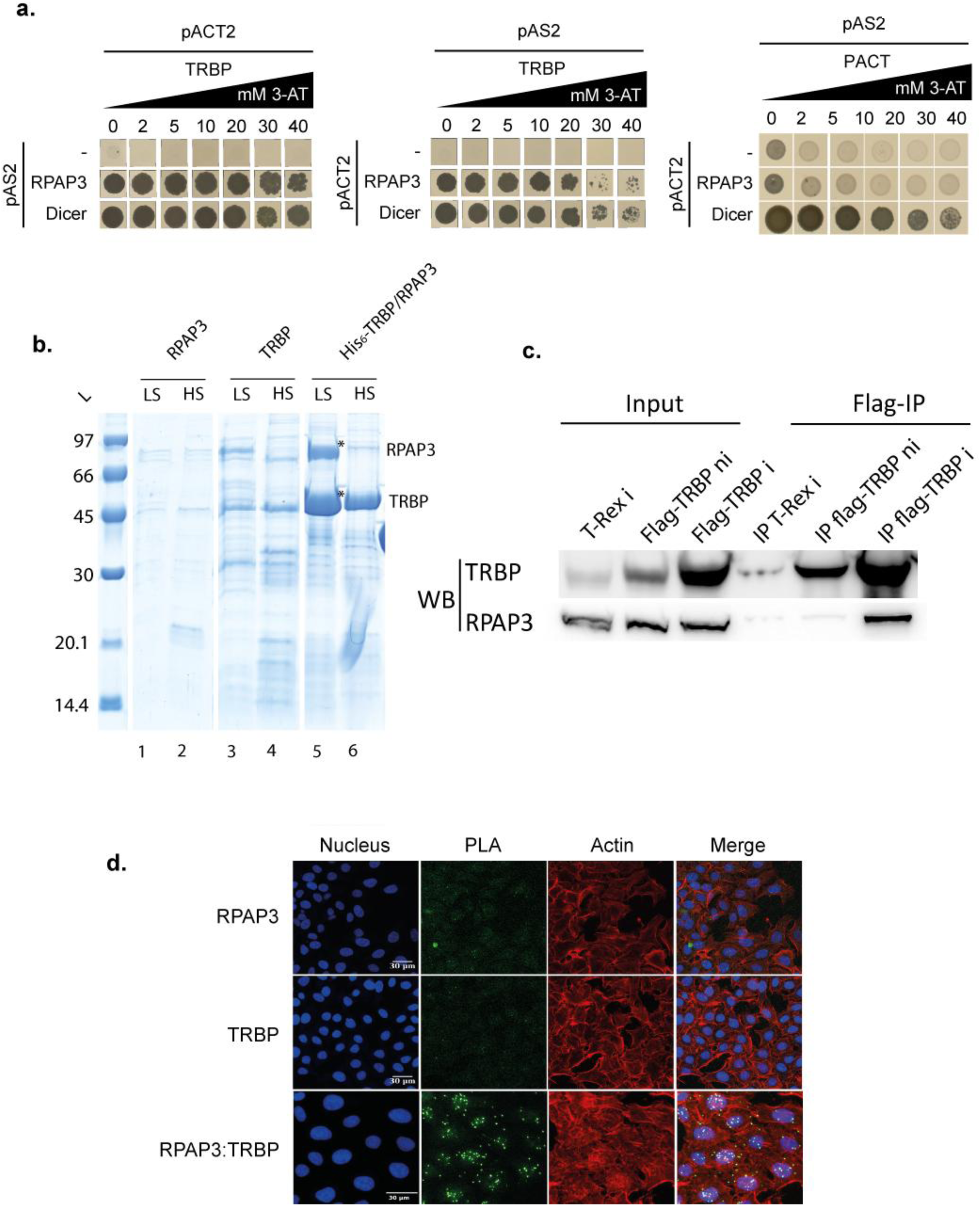
Characterization of the TRBP:RPAP3 interaction. **a**. A candidate-based yeast two-hybrid (Y2H) screen performed in the yeast *S. cerevisiae* revealed an interaction between TRBP and RPAP3 in both orientations (left and middle panels). On the contrary, RPAP3 does not bind to PACT (right panel). The TRBP:Dicer interaction was used as a positive control. pAS2 and pACT2 plasmids expressed a protein fusion with the DNA binding domain or activation domain of transcription factor Gal4, respectively. The strength of the interactions was tested using increasing amounts of 3-amino-triazol (3-AT) from 0 to 40 mM. **b**. Co-expression experiments in *E. coli*. RPAP3 copurified with a hexahistidine His_6_-tagged version of TRBP on cobalt-based immobilized metal affinity chromatography (IMAC) beads (TALON) at low salt concentration (LS), but not in high-salt conditions (HS). Single protein expression controls experiments for both untagged TRBP or RPAP3 proteins are shown in lanes 1-4. See materials and methods for details. **c**. Co-immunoprecipitation experiments performed in the T-Rex HEK293 cell line expressing a flagged version of TRBP upon doxycycline induction. ni: no doxycycline induction, i: doxycycline induction. RPAP3 is co-immunoprecipitated with the flagged TRBP protein. **d**. *In cellulo* Duolink assays performed in HeLa cells. Proximity ligation assay (PLA) reveals a close proximity of the endogenous TRBP and RPAP3 proteins in favor of their direct interaction. Nuclei were stained using DAPI, and cytoplasmatic actin using Alexa Fluor 546. Scale bar is 30 μm. See Materials and Methods for details.

### The TPR1 domain of RPAP3 binds the non-canonical type B dsRBD (dsRBD3) of TRBP

To define the domain of RPAP3 that mediates the interaction with TRBP, we performed co-expression experiments in *E. coli* and Y2H assays, using different protein sub-domains **(Fig. 2, Supp. Fig. 4)**. In metazoan, RPAP3 contains two tetratricopeptide (TPR) domains, each composed of 3 TPR motifs and a capping helix at the C-terminal end (39,43) **(Fig. 2 a)**. Two highly soluble TPR domains have been defined in RPAP3, TPR1 encompassing residues 133 to 255 and TPR2 including residues 281 to 396. On the other hand, TRBP folds into 3 evolutionary conserved dsRBDs (11). The two N-terminal ones mediate dsRNAs binding (residues 18-99 and 157-228, respectively, 44,45), while the C-terminal one mediates the interaction with Dicer (residues 262-366; **Fig. 2 a)**. By using co-expression assay in *E. coli* **(Fig. 2 b)** and yeast two hybrids assay **(Fig. 2 c left, Supp. Fig. 4 b left)**, we found that the TPR1 domain of RPAP3 is both necessary and sufficient for TRBP binding. Surprisingly, this is not the case for the TRP2 domain, in spite of its strong homology with TPR1. Next, we used a similar strategy to define the region of TRBP required for RPAP3 binding. We found that the dsRBD3 domain (262-366) of TRBP **(Fig. 2 d right panel, Supp. Fig. 4 b)**, but not the dsRBD1 and dsRBD2 domains **(Fig. 2 d left and middle panels, Supp. Fig. 4 b)**, interacted with the TPR1 domain of RPAP3, which was confirmed using Y2H assay **(Fig. 2 c right, Supp. Fig. 4 a right)**.

**Figure 2.**
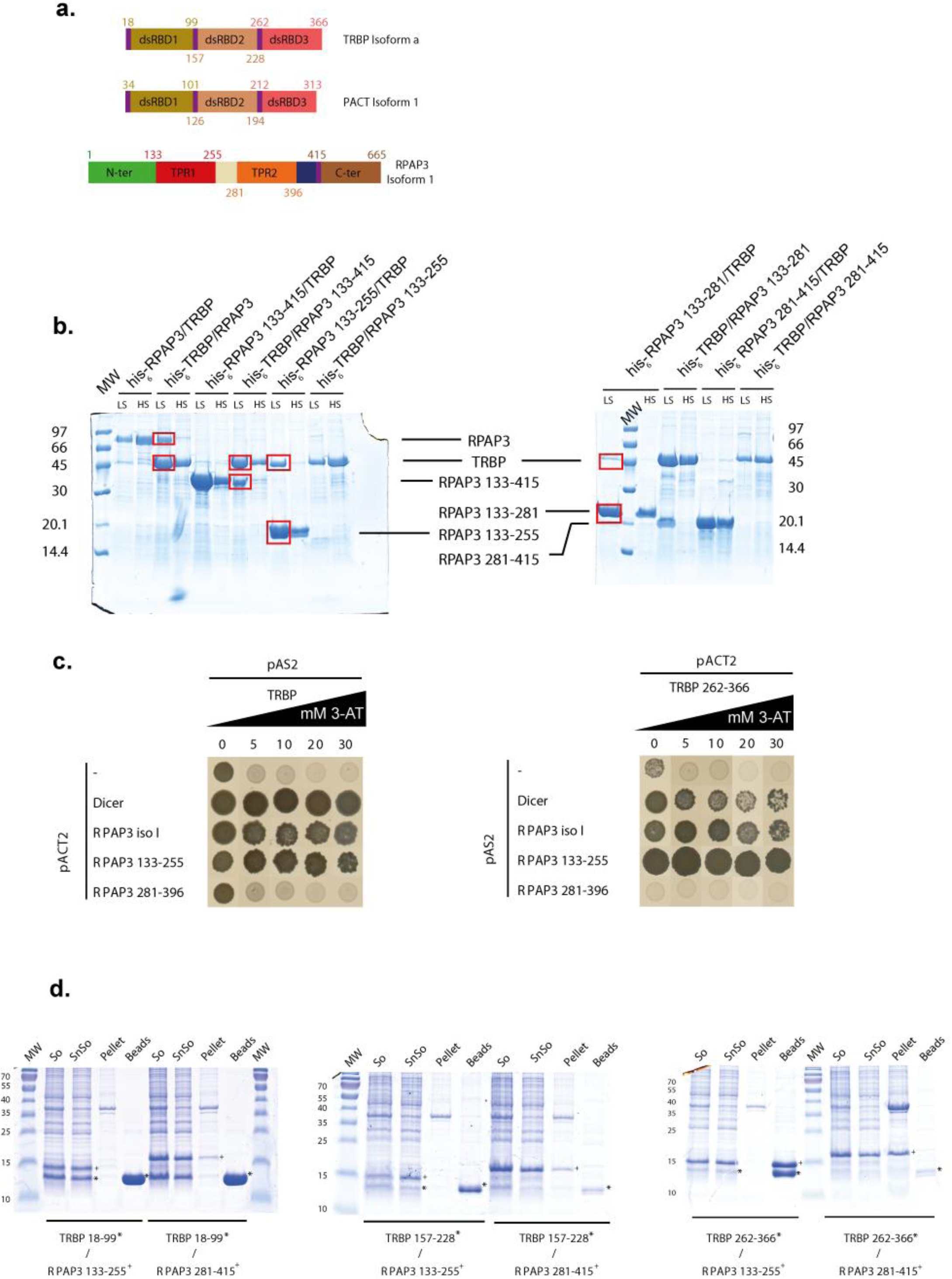
Identification of the protein subdomains involved in the interaction. **a**. Representation of the subdomains of TRBP and RPAP3 as predicted with the software IUPRED and PSIPRED (69,70). Numbers correspond to the amino-acids residues located at each domain extremities. **b**. Co-expression experiments in *E. coli* performed with RPAP3, TRBP and their subdomains. For each protein pair, the first protein name on top of the gel always indicates the His_6_-tagged protein. Positive interactions are highlighted with red squares. **C** Two-hybrid screens performed in the yeast *S. cerevisiae* on TRBP, RPAP3 and their subdomains. The TRBP:Dicer interaction was used as a positive control. pAS2 and pACT2 plasmids respectively enable expression of a protein fusion with the DNA binding domain or activation domain of transcription factor Gal4. Strengths of the interactions were tested using increasing amounts of 3-amino-triazol (3-AT) from 0 to 40 mM. **D**. Co-expression experiments in *E. coli* performed with RPAP3 and TRBP subdomains. The TRBP and RPAP3 domains used are indicated below the gel lanes. The TRBP subdomains carried the His_6_-tag. “So”, “SnSo”, “Beads”, “MW” and “Pellet” design respectively the culture sonicate, supernatant sonicate, Talon beads, Molecular Weight marker and sonicate pellet.

Finally, to define more precisely the RPAP3-binding site in the TRBP dsRBD3, we performed similar experiments with shorter TRBP fragments **(Supp. Fig. 4 c)**. Collectively, these experiments revealed that the TRBP dsRBD3 C-terminal part, residues 293-366, was sufficient for RPAP3 binding. We concluded that the RPAP3 TPR1 domain interacts directly with the TRBP dsRBD3, and more precisely with its residues 293 to 366.

### The TRBP:RPAP3 and the TRBP:Dicer interactions are mutually exclusive

The dsRBD3 domain of TRBP was shown to be the domain involved in the interaction with Dicer (10,11,44,46). Therefore, to assess whether a ternary complex containing Dicer, TRBP and RPAP3 could be formed, we performed simultaneous co-expression experiments in *E. coli* of the three protein minimal subdomains involved in the respective interactions **(Fig. 3 a-c)**. We found that both the TPR1 domain of RPAP3 and the 256-595 domain of Dicer co-purified with a His_6_-tagged version of TRBP dsRBD3 on TALON^®^ beads **(Fig. 3 a)**. However, when the eluate was loaded onto a gel filtration column, the elution profile clearly showed two distinct peaks corresponding to TRBP:RPAP3 and TRBP:Dicer sub-complexes, respectively **(Fig. 3 b, c, Supp. Fig. 4 d)**. This indicated that the dsRBD3 of TRBP could not interact simultaneously with Dicer and RPAP3, probably due to steric constrains as the same domain of TRBP seems to be involved in both interactions.

**Figure 3.**
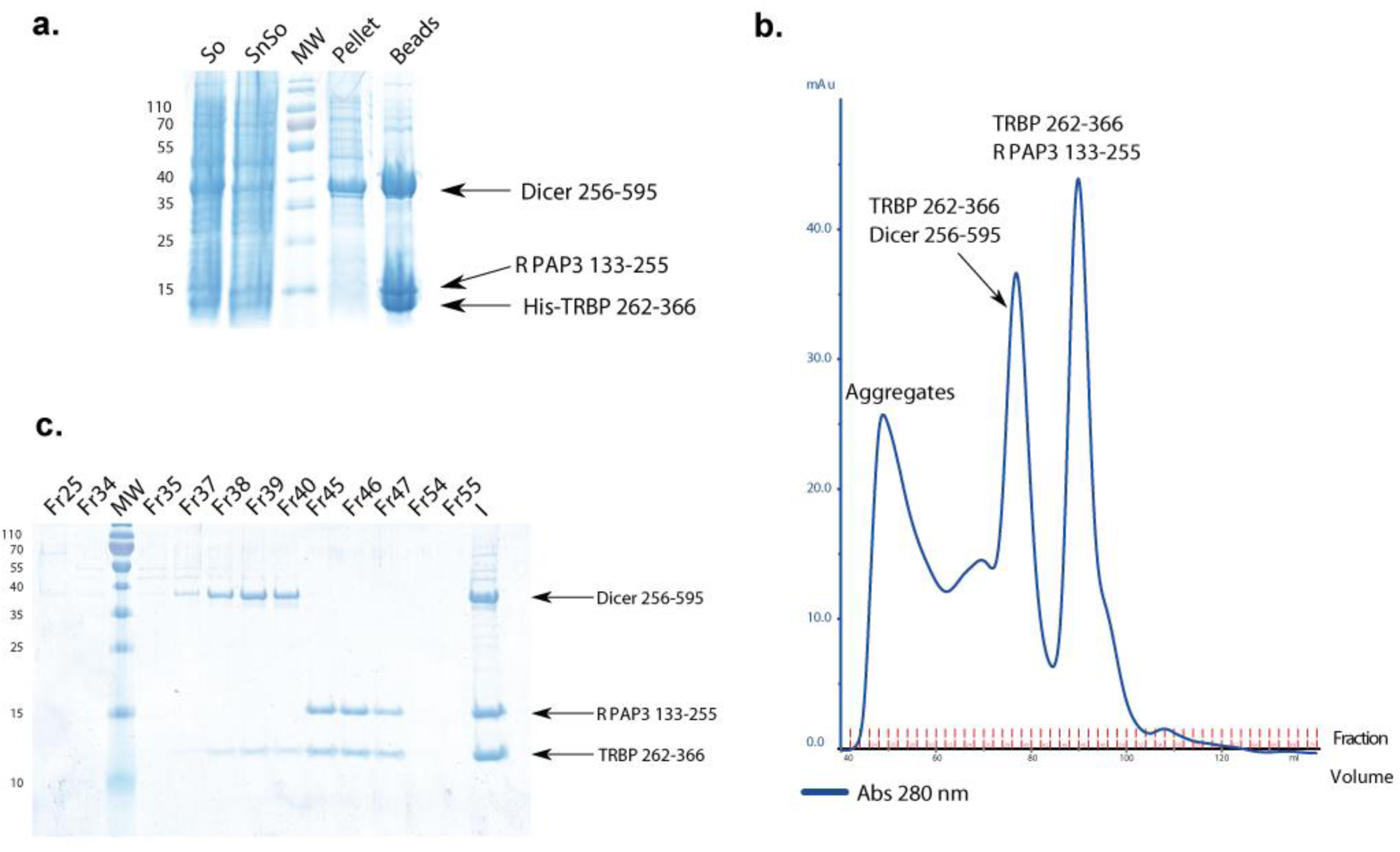
The same binding surface of TRBP seems to be involved for its interactions with Dicer or RPAP3. **a**. Co-expression experiments in *E. coli* of the three minimal protein subdomains involved in the TRBP:RPAP3 and TRBP:Dicer interactions. “So”, “SnSo”, “Beads”, “MW” and “Pellet” relate respectively to the culture sonicate, supernatant sonicate, Talon beads, Molecular Weight markers and sonicate pellet. **b**. Elution profile of the gel filtration assay performed on the 3 co-expressed TRBP, RPAP3 and TRBP minimal subdomains. **c**. Fractions collected were loaded onto a 10% SDS-PAGE.

### Structure of the human RPAP3:TRBP complex

In order to highlight the structural features of the interaction between the TPR1 domain of RPAP3 and the dsRBD3 of TRBP, we crystallized the heterodimer and collected X-ray diffraction data at 1.49 Å resolution **(Fig. 4, Table 1;** see Materials and Methods). The RPAP3 core (residues 133-249) consists of seven α-helices arranged in a repeating antiparallel right-handed helix topology and was already described as a TPR (tetratricopeptide repeat) domain in the crystal structure of RPAP3 bound to the C-terminal tail peptide (SRMEEVD) of HSP90 (35,37,39) **(Supp. Fig. 5 a, in blue)**. On the other hand, the TRBP structure (residues 262-366) contains a α/β sandwich (residues 289-366) typical of a dsRBD fold **(Supp. Fig. 5 b, in purple)**, with three β-strands (β1,β2, and β3) and two α-helices (H4 and H5), as already described (10). Interestingly, the crystal structure of RPAP3 (residues 133-249) bound to TRBP is similar to that of RPAP3 bound to the C-terminal tail peptide (SRMEEVD) of HSP90 (35,37,39) **Supp. Fig. 5 a, in green)**. Similarly, the dsRBD core of TRBP bound to RPAP3 is structurally similar to the dsRBD domain of TRBP in complex with Dicer (10) **(Supp. Fig. 5 b, in orange)**. These observations suggest that no significant conformational modification occurs on neither protein upon binding to each other. Interestingly, however, an N-terminal extension (residues 262-288) beyond the canonical dsRBD domain of TRBP is observed in the crystal structure of the RPAP3:TRBP complex and consists of 3 helices H1, H2 and H3 **(Supp. Fig. 5 b, in orange, and Supp. Fig. 5 c)**. This N-terminal extension was shown to be partially disordered in the crystal structure of TRBP bound to Dicer (10) (see supplementary materials for a detailed structural description).

**Figure 4.**
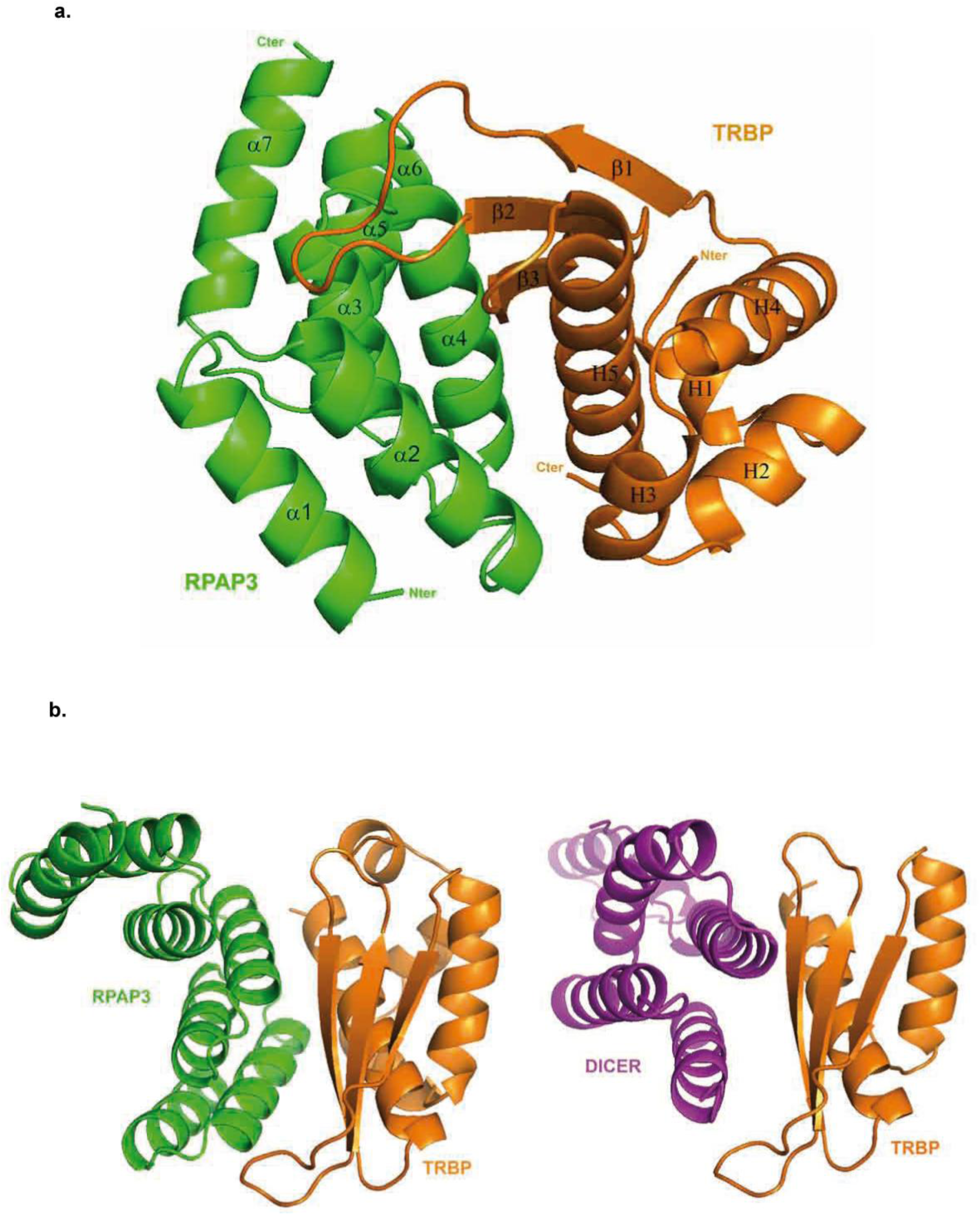
Crystal structures of the TRBP:RPAP3 complex. **a**. Ribbon representation of the X-ray crystallographic structure of the TPR1 domain of RPAP3 and the dsRBD3 of TRBP at 1.49 Å resolution. RPAP3 (residues 133-249) and TRBP (residues 263-365) are drawn in green and orange, respectively. **b**. Ribbon representations of our crystal structure (left) and the one published by Wilson *et* al. (23) (right) confirm that TRBP shares the same binding interface for its interaction with either Dicer or RPAP3.

As suspected from our co-expression and gel filtration experimental results **(Fig. 3)**, the crystal structure revealed that the protein interface involves the second α-helix H5 as well as β-strands β2 and β3 of the dsRBD3 of TRBP **(Fig. 4 a)**. We confirmed this data using solution-state NMR spectroscopy. We assigned backbone resonances of the RPAP3-TPR1:TRPB-dsRBD3 complex and, thanks to TALOS-N calculations, we showed that the two partners fold similarly in solution and in the crystal (**Supp. Fig. 6**). Then, we measured chemical shift perturbations of backbone amide groups in RPAP3-TPR1 upon binding of TRBP-dsRBD3 **(Fig. 5 a, b)**. This NMR mapping revealed that major perturbations are observed in helices α2 and α4 of RPAP3-TPR1, in accordance with the binding interface observed in the X-ray structure. Since these helices are also involved in the interaction with Dicer (10), this confirmed that the TRBP interaction with RPAP3 or Dicer are mutually exclusive in solution, even if Dicer and RPAP3 do not share any overall structure similarities **(Fig. 4 b)**.

**Figure 5.**
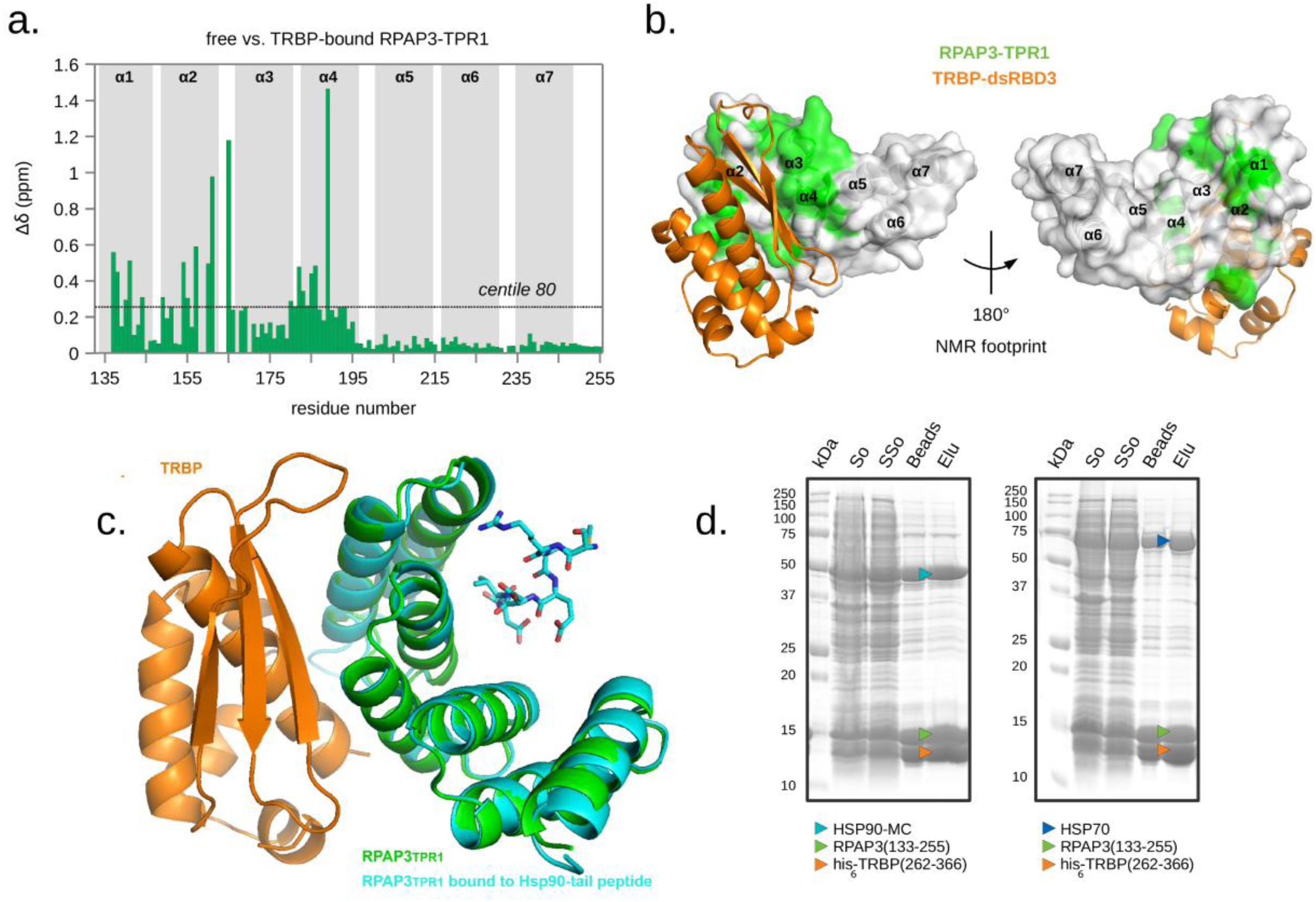
Two distinct surfaces of RPAP3 bind TRBP and the HSP90-tail peptide. **a**. Chemical shift perturbations of RPAP3-TPR1 upon binding of TRBP-dsRBD3. Backbone amide group resonances of the free and TRPB bound-states of RPAP3-TPR1 were compared using composite ^1^H-^15^N chemical shifts (Δδ). Data were plotted againt the sequence of RPAP3. Position of helices was indicated in grey and the value corresponding to centile 80 was indicated with a dotted line. **b**. Residues in RPAP3 for which the Δδ value was superior to the centile 80 value were reported in green on the molecular surface of RPAP3-TPR1. The X-ray structure of the RPAP3-TPR1:TRBP-dsRBD3 complex was used. **c**. Superimposition of the crystal structure of RPAP3-TRBP (in green and orange) with the crystal structure of RPAP3 bound to the HSP90-tail peptide (in cyan) (30) (PDB 4cgv). The HSP90-tail peptide (SRMEEVD) is shown as sticks. **d**. Protein co-expression assays in *E. coli* with the His_6_-TRBP-RPAP3 complex and human HSP70 or HSP90-MC. “So”, “SSo”, “Beads” and “Elu” relate respectively to the culture sonicate, supernatant sonicate, Talon beads and elution from the beads with imidazole. The co-purified proteins are indicated with colored arrows.

In contrast, binding of RPAP3 to TRBP involves the convex surface, i.e. the opposite face of the TPR domain compared to that involved in HSP90 binding, suggesting that HSP90 binding should not be altered by TRBP binding (a detailed structural description is available in the supplementary materials section). Superimposition of the RPAP3:TRBP structure with the structure of RPAP3 (TPR1) bound to the C-terminal tail peptide (SRMEEVD) of HSP90 (35,37) indeed shows that the RPAP3 surface binding to the HSP90-tail peptide is far away from the RPAP3:TRBP interface **(Fig. 5 c)**. We concluded that binding of TRBP to RPAP3 should not prevent the recruitment of HSP90 and HSP70 by the R2TP complex. This prediction was experimentally verified by co-expression experiments in *E. coli* between different domains of RPAP3 and HSP70/90, and TRBP (262-366) **(Fig. 5 d**., **Supp. Fig. 7)**.

### Identification of key residues involved in the interaction between RPAP3 and TRBP

Based on the crystal structure of the interface between human TRBP and RPAP3, we performed a mutational analysis and tested the TRBP:RPAP3 interaction by co-expression in *E. coli* and coimmunoprecipitation assays. We substituted several residues within the TPR1 domain of RPAP3 with alanine **(Supp. Table 1)**. Some of the mutated proteins, e.g. L192A **(Supp. Fig. 8 a)**, were expressed at low levels in *E. coli*, suggesting that the mutations affected the folding and/or stability/solubility of the protein (data not shown). Then, we investigated the conserved inter-protein polar contacts involving side-chains and identified the RPAP3:TRBP residues interactions between D150 (RPAP3) and S320 (TRBP), T157 (RPAP3) and R354 (TRBP) or D161 (RPAP3) and Q357 (TRBP) (hydrogen bonds) as important **(Fig. 6, Supp. Fig. 8)**. In agreement with structural data, individual mutations of all these residues except for S320A in TRBP, destabilized the RPAP3:TRBP interaction. Interestingly, the point mutation V185A on RPAP3 had a drastic effect on TRBP binding without affecting protein solubility, showing that this residue is crucial for the interaction as assumed from the structure of the complex **(Supp. Fig. 8 b)**. Next, we tested the interactions in human cells using IP-LUMIER experiments. The mutation of D161A on RPAP3, which was found hydrogen bonded to Q357 in TRBP, disrupted the complex **(Fig. 6)**. This pair of residues was particularly interesting as mutant Q357A in TRBP also disrupted its interaction with RPAP3, but not with Dicer, as revealed by IP-LUMIER experiments **(Fig. 6 b, c)**. This mutagenesis analysis thus identifies a residue of TRBP whose mutation selectively affects binding to RPAP3 but not Dicer. Interestingly, this D161 amino acid is substituted by A310 in the TPR2 of RPAP3, which could explain why TRBP does not bind this second TPR domain.

**Figure 6.**
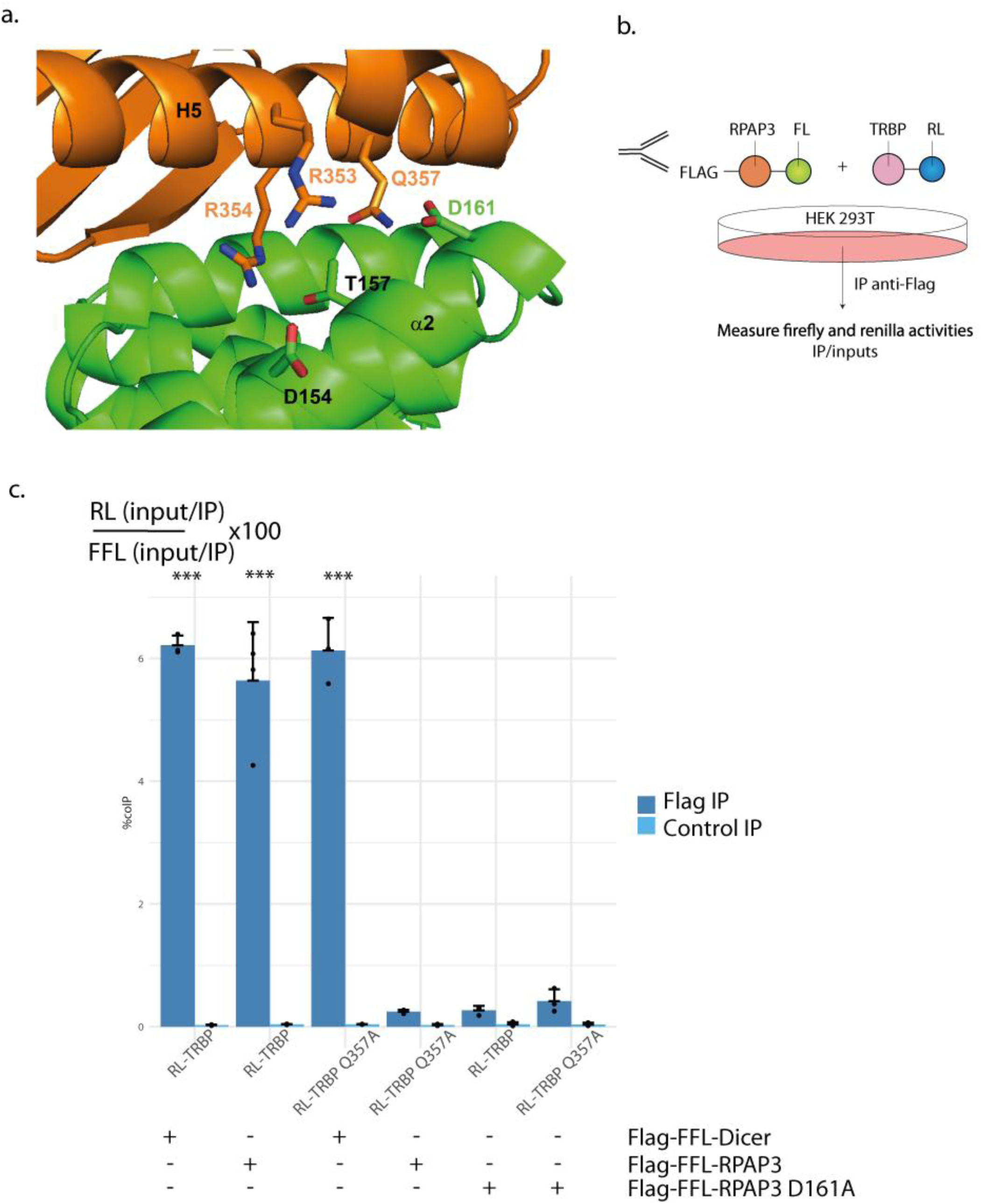
Identification of key residues involved in TRBP:RPAP3 binding interface. **a**. Network of ionic interactions as well as hydrogen bonds between RPAP3 and TRBP. RPAP3 and TRBP are drawn in green and orange, respectively. **b**. Principle of the LUMIER-IP test. **c**. Bar plot showing LUMIER-IP co-efficiency for the interaction of Dicer with TRBP WT or Q357A. *** p-value <0,001 (Z-test comparing values of the FLAG IPs with eleven-times the mean value obtained in the control IP) or the interaction of RPAP3 D161A with TRBP WT or Q357A.

### TRBP, Dicer and AGOs require HSP90 activity

We showed that TRBP, RPAP3 and HSP90 could form a ternary complex together **(Fig. 5)**. In order to investigate the possible involvement of the entire R2TP complex rather than RPAP3 alone, we verified whether TRBP was able to coprecipitate R2TP core proteins other than RPAP3, using a transiently expressed V5 tagged-TRBP. We observed that all the R2TP proteins, namely PIH1D1 and RUVBL1/2 were efficiently coprecipitated by V5-TRBP **(Fig. 7 a, lane 3)**. This is an interesting result, as we did not detect TRBP interactions with PIH1D1 or the RUVBL1/2 proteins in our initial candidate-based yeast two-hybrid screen. This may be explained by an indirect interaction mediated by RPAP3, and indeed most interactions were lost when we used TRBP mutant R354E, defective for RPAP3 binding **(Fig. 7 a, lane 1)**. However, even when the interaction with RPAP3 is lacking, TRBP could still co-precipitate RUVBL2 **(Fig. 7 a, lane 1)**, indicating a connection between TRBP and RUVBL2.

**Figure 7.**
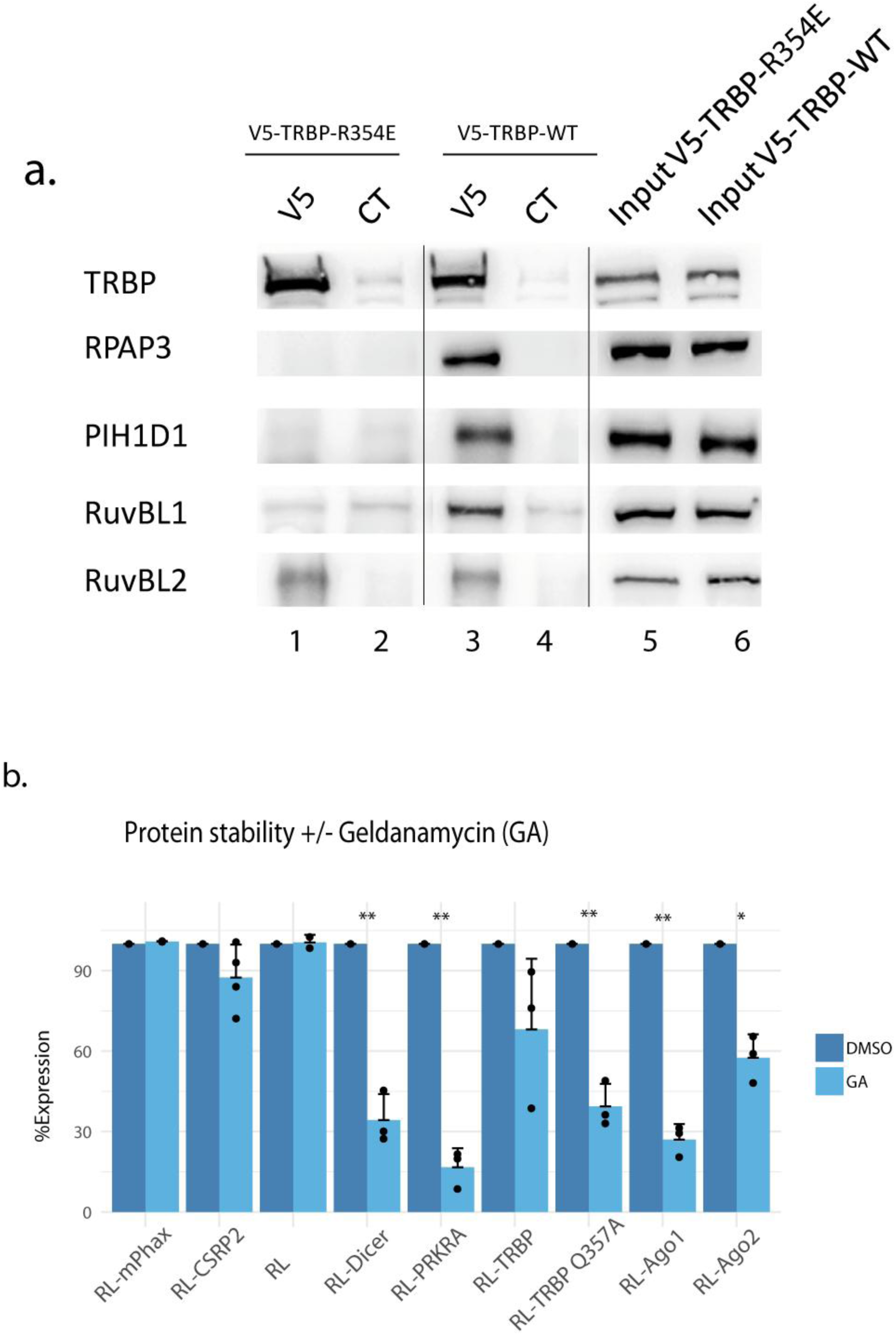
TRBP and Dicer are functionally linked to the R2TP/HSP90 complex. **a**. Co-immunoprecipitation experiments performed using a transiently expressed V5-tagged TRBP protein, or the TRBP R354E. RPAP3, PIH1D1, RUVBL1 and RUVBL2 are the components of the R2TP complex in human. **b**. Bar plots representing the ratio RL/FFL to show the stability of RL-Dicer, RL-AGO1/2 and RL-TRBP WT/Q357A in 293T cell treated with DMSO or DMSO+ 2 µM of geldanamycin during 16 h. Plasmid expressing FFL alone is used to normalize the value obtained for RL protein in each assay. Phax and CSRP2 are two unrelated protein used as negative controls. * p-value <0,05. ** p-value <0,01 according to a Student test. % expression = 100*(RL-X_GA_/FFL_GA_) / (RL-X_DMSO_/FFL_DMSO_).

Finally, in order to determine whether TRBP, Dicer or AGO1/2 proteins could be clients of the HSP90/R2TP chaperoning system, we tested their stability after HSP90 inhibition with geldanamycin, a drug often leading to HSP90 client destabilization (22,26). To this end, we transfected plasmids expressing each of these proteins fused to the Renilla luciferase together with a Firefly luciferase control vector **(Fig. 7 b)**. Remarkably, all proteins were sensitive to Geldanamycin, revealing the importance of HSP90 for their stability. Interestingly, the mutant of TRBP that does not bind RPAP3 (TRBP Q357A) seems to be more affected by geldanamycin, which suggests a stabilizing role for RPAP3 on TRBP.

## DISCUSSION

### RPAP3 binds to TRBP using the same surface as Dicer

Our work identified a yet undescribed direct interaction between RPAP3, a core component of the HSP90/R2TP chaperone system, and TRBP, which, among other functions, is one of two alternative Dicer cofactors. We also showed that RPAP3 and Dicer use the same surface of TRBP for binding and thus that the two interactions were mutually exclusive **(Fig. 4)**. This was surprising, as there is no significant similarity between RPAP3 and Dicer, neither at the sequence, nor at the secondary structure, levels. For deeper comparison, we superimposed the RPAP3:TRBP and Dicer:TRBP 3D structures using only atoms from TRBP. As expected, the overall 3D structures of RPAP3 and Dicer do not superimpose to each other and the superimposition revealed no significant conformational modifications of TRBP whether it binds to RPAP3 or to Dicer **(Supp. Fig. 5)**. However, a detailed analysis of both interactions revealed that the central parts of α-helixes α4 from RPAP3 and Dicer are located at the same position on the surface of TRBP **(Supp. Fig. 9 a, b)**. Thus, despite very different overall 3D structures, RPAP3 and Dicer share a similar binding site at the TRBP surface and display a α-helix (named α4 in both proteins) that allows equivalent interactions in both RPAP3:TRBP and Dicer:TRBP complexes. This helix could thus be the key determinant for other uninvestigated protein recruitment by TRBP.

### RPAP3 does not interact with PACT unlike Dicer

Dicer can associate with either TRBP or PACT and the Dicer:TRBP and Dicer:PACT complexes selectively contribute to miRNA length and strand selection in mammalian cells. Indeed, TRBP and PACT differentially affect dsRNA structure and orientation on Dicer, resulting in different Dicer pre-miRNAs processing activities (10,12). PACT and TRBP are paralogs and their structural organization are very similar. Indeed, TRBP and PACT bind Dicer in a similar manner and their interactions are mutually exclusive (10). As shown above, RPAP3 share some common structural features with Dicer that allow the binding to TRBP. Our candidate-based yeast-two hybrid screen **(Fig. 1a)** completed by co-expression assays in *E. coli* revealed no interaction between RPAP3 and PACT **(Supp. Fig. 1 e-f)**. This is surprising based on the high amino-acid sequence identity of the third dsRBD of PACT and the homologous sequence in TRBP (55%). Sequence alignment **(Supp. Fig. 9 c)** reveals that six over nine residues involved in the RPAP3:TRBP interface are strictly conserved between PACT and TRBP. Only residues S320, R353 and R354 in TRBP are substituted for N, H and N in PACT, respectively. As shown above in the RPAP3:TRBP complex, the side-chain of S320 of TRBP forms a hydrogen-bond with the side-chain of D150 in RPAP3 **(Supp. Fig. 8 c)**. Substitution of S320 in TRBP for N in PACT does not abolish possible hydrogen-bond formation. On the other hand, the positively-charged residues R353 and R354 forming ionic interactions at the RPAP3:TRBP interface **(Fig. 6 a)** are also respectively substituted for H and N in PACT, abolishing the possibility to form salt-bridges. Moreover, residue R354 in TRBP is crucial for binding to both RPAP3 and Dicer. Indeed, the R354E mutation disrupts both complexes. Noticeably, the TRBP variant Q357A still interacts with Dicer while it no longer binds RPAP3. Thus, Dicer can bind wild-type TRBP, TRBP Q357A, but not TRBP R354E. However, Dicer is able to bind to PACT where R354 is substituted for N as compared to TRBP. On the other hand, RPAP3 binds wild-type TRBP but not the variants R354E, Q357A and does not bind PACT at all. Altogether, this suggests that substitution of R354 for N in PACT (N301) may be a key point to explain why TRBP binds RPAP3 but PACT does not. Another explanation could be the fact that PACT homodimerizes through its dsRBD3 more strongly than TRBP does, which we observed by NMR and native mass spectrometry (data not shown), thus preventing RPAP3 binding (44).

The mutually exclusive interaction we have described here, between TRBP and RPAP3 or between TRBP and Dicer, associated to the absence of interaction between RPAP3 and PACT, could reveal a putatively novel regulation mechanism of Dicer activity. Indeed, the final processing step of pre-miRNAs by Dicer in the cytoplasm is crucial to generate the proper miRNA ends and thus to specify its mRNA binding properties. PACT and TRBP, the two Dicer partners, have distinct effects on Dicer-mediated dsRNA processing (10,13,45). RNAs processed from long dsRNAs are for example not loaded by PACT, while TRBP handles both pre-miRNA and long dsRNAs. It was also shown that Dicer differentially generated isomiRs depending on its association to TRBP, PACT or none of them (10). Regulation of TRBP binding to Dicer by sequestration via RPAP3 could therefore introduce new biases in miRNA processing, and subsequently mRNA target specificity.

### TRBP and Dicer stabilities rely on HSP90

We showed that RPAP3 binds HSC70/HSP90 chaperones when associated to TRBP **(Fig. 5 a, b, Supp. Fig. 7)**. HSC70/HSP90 dependent RISC loading could therefore involve RPAP3 to some extent (15–17,47,48), **Fig. 8)**. We also showed that TRBP, probably via RPAP3, co-precipitated all members of the R2TP complex, and that the interactions between TRBP and RPAP3, and between TRBP and Dicer, were mutually exclusive. This could reflect that RPAP3 is involved in the TRBP:Dicer assembly, or recycling of either proteins following pre-miRNA cleavage. Indeed, the mutually exclusive character of the TRBP:RPAP3 or TRBP:Dicer interactions reflects a possible exchange mechanisms, where RPAP3 would be replaced by Dicer on TRBP, or *vice versa* **(Fig. 8, *1*.)**. TRBP and Dicer could therefore constitute new clients for the R2TP complex **(Fig. 8)**.

**Figure 8.**
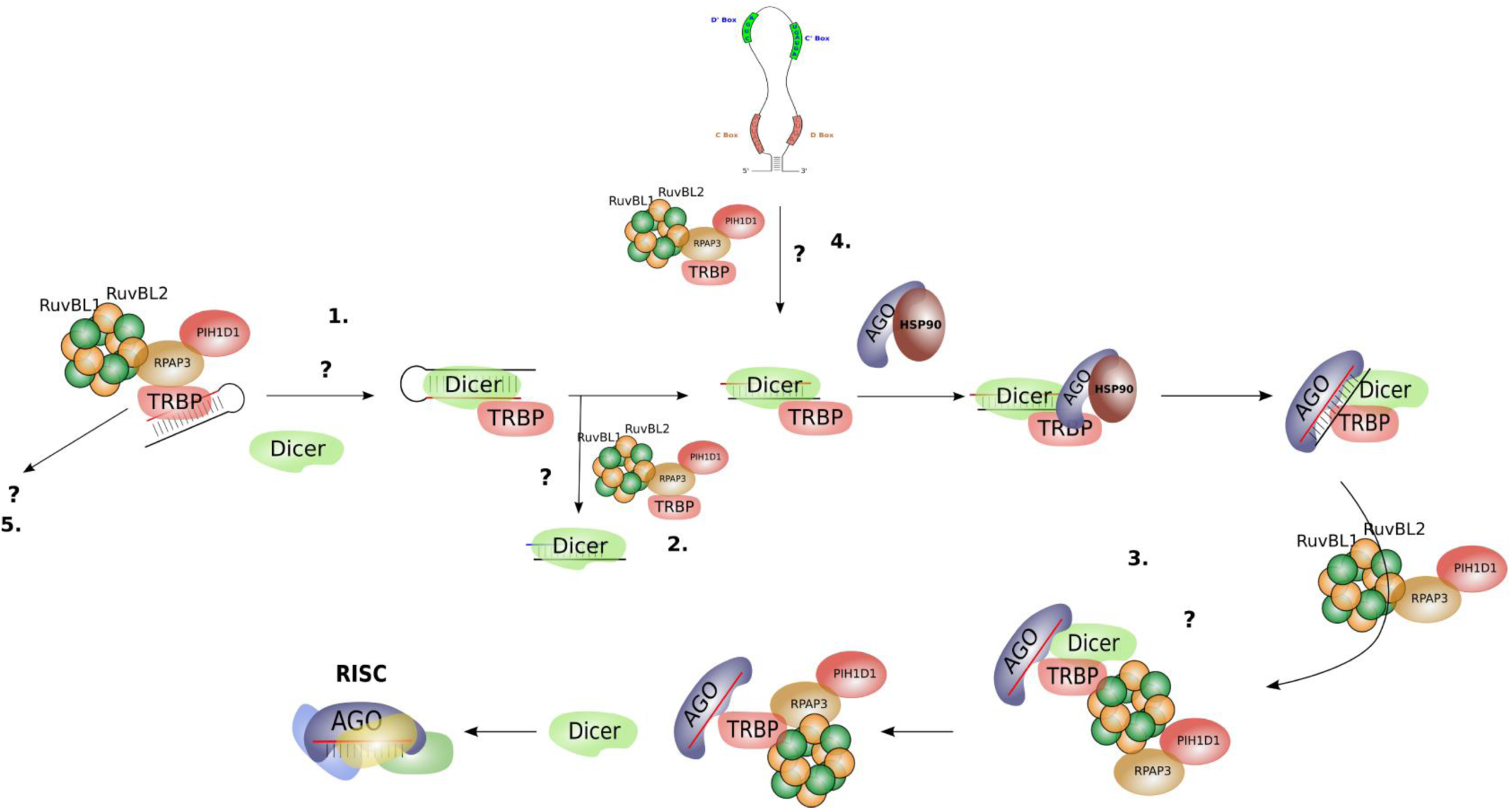
Hypothetical functions for the TRBP:RPAP3 interaction. See text for details.

Another possibility could be that RPAP3 and/or the R2TP complex provides some specificity to the RNAs cleaved by TRBP:Dicer **(Fig. 8, *2*.)** or loaded onto the RISC **(Fig. 8, *3*.)**. Indeed, alternative maturation pathways of miRNA have been described (49,50), and a particularly interesting possibility consists in the miRNAs or miRNA-like RNAs that are processed from small nucleolar RNAs (snoRNAs). These miRNAs are referred to as sno-derived RNAs or sdRNAs (51–55), and their maturation pathways follow poorly understood mechanisms. However, as the R2TP is well known to be involved in canonical snoRNP assembly, it provides a molecular link between snoRNAs and miRNAs, and it is tempting to speculate that it may be involved in the generation of sdRNAs **(Fig. 8, *4*.)** (22,51,56). Interestingly, AGO proteins or DGCR8 constitute another link between the two process as both are known for their roles in miRNA functions, and as they were shown to be associated to numerous snoRNAs (51,57).

Finally, RPAP3 could be involved in Dicer-independent additional functions of TRBP. **(Fig. 8, *5*.)**. For instance, RPAP3 could simply stabilize neo-synthesized TRBP, before binding to its final partner, whatever it is, and which function this partner plays. Indeed, we showed that TRBP is normally weakly sensitive to HSP90 inhibition, but the TRBP Q357A mutant that does not bind RPAP3 anymore becomes much more sensitive to HSP90 activity, possibly because the mutation, and the absence of RPAP3 binding, destabilizes TRBP so that it now requires HSP90. RPAP3 would here simply play a role of free TRBP stabilization, before or after its interaction with Dicer for example.

## METHODS

### Cell culture

HeLa and HEK293T (including T-Rex cell lines) cells were maintained in Dulbecco’s modified Eagle’s medium. HCT116 cells were maintained in Mc’Coys medium. Both media were supplemented with 10% of fetal bovine serum, 10 U/ml of penicillin/streptomycin and 2.9 mg/ml of glutamine, in a humidified 5% CO_2_ incubator at 37°C. Additionally, T-Rex cells were maintained with 100 µg/ml of zeocin 10 µg/ml of blasticidin.

### Generation of stable, inducible, Flag T-REX cell lines

Inducible T-Rex cell lines were generated following the manufacturer’s instructions (Invitrogen). Briefly, confluent HEK-293 T-Rex cells were transfected in 10 cm cell culture plates in blasticidin containing medium (no zeocin) with 9 µg of the pOG44 plasmid enabling expression of the Flp recombinase and 1 µg of the pcDNA5/FRT plasmid containing the flagged protein of interest gene (RPAP3 or TRBP). Medium was changed after two days with blasticidin containing medium. The next day; cells were split and treated with 100 µg/ml of hygromycin B. The blasticidin/hygromycin medium is changed every 4 to 5 days, during two weeks, until isolated clones can be retrieved and transferred to a new dish for screening.

### Plasmids and cloning

DNA cloning was performed using standard techniques or with Gateway™ system (Invitrogen). For NMR and crystallogenesis assays, TRBP and RPAP3 ORFs were cloned in pnEA-3CH and pnCS vector (respectively) at the *5’-NdeI* and *3’-BamHI* sites (58,59). For co-expression assays, RPAP3, TRBP, Dicer and PACT ORFs were cloned in pnEA-3CH, pnCS and pnYK plasmids modified to become compatible with the Gateway cloning technology. For that, the *ccdb* and *chloramphenicol* genes were amplified by PCR in pDEST17 and inserted in these vectors at the *5’-NdeI* and *3’-BamHI* sites. pnYK vector was created from pnYC vector (58) by homologous recombination between the chloramphenicol and kanamycin genes resistance gene using the In-Fusion kit (Clontech) following the manufacturer’s recommendations. For GST pull-down experiments, ORFs were cloned in pDEST15 (containing the GST tag) and pDEST17 Gateway (containing the His_6_ tag) vectors (Thermo Fisher Scientific). For Y2H assays, pACT2, pAS2 and pGBKT7 were used. For co-immunoprecipitation, the TRBP ORF was cloned into the pcDNA3.1 and nV5-DEST plasmids (Invitrogen). For the LUMIER-IP and Luciferase assays, pcDNA5-FRT-3xFLAG-FFL-Rf (CMV promoter), and L30-HA-RL were used. The cDNAs were all of human origin except for PHAX which was from mouse.

### Antibodies

Antibodies and dilutions for IF and Duolink were the following: mouse monoclonal anti-RPAP3 at 1:250 dilution (Sigma-Aldrich, SAB407956); polyclonal rabbit anti-TRBP at 1:100 dilution (Abcam, ab72110); monoclonal mouse anti-Actin at 1:400 dilution (Abcam, ab3280); polyclonal rabbit anti-GAPDH at 1:750 dilution (Abcam, ab9485); monoclonal mouse anti-GAPDH at 1:750 dilution (Abcam, ab8245). Antibodies and dilutions for western-blot were the following: rabbit polyclonal anti-RPAP3 at 1:2000 dilution (Sigma-Aldrich, SAB1411438); monoclonal mouse anti-TRBP at 1:500 dilution (Abcam, ab129325); polyclonal rabbit anti-TRBP at 1:500 dilution (Abcam, ab72110); polyclonal rabbit anti-V5 at 1:4000 dilution (ThermoFischer Scientific, GTX117997); antibodies for IP was the following monoclonal mouse anti-V5 (ThermoFischer Scientific, 37-7500).

### Yeast Two Hybrid (Y2H)

For Y2H assays, appropriate pACT2 and pAS2 plasmids were introduced into haploid *S. cerevisiae* test strains (Y187 and CG1945, respectively), which were then crossed. Diploids were selected on Leu^-^/Trp^-^ medium and then plated on Leu^-^/Trp^-^/^His-^ plates, with 0 to 40 mM of 3-Amino-1,2,4-Triazol (3-AT), which is a competitive inhibitor of the product of the *HIS3* reporter gene. This was used to evaluate the strength of the interactions. Growth was assessed after three or four days of incubation at 30°C (60).

### Co-expression experiments in *E. coli*, protein production and purification

For crystallogenesis assays, *E. coli* BL21 (DE3) pRARE2 cells were co-transformed with the non gateway version of the pnEA3CH::TRBP (266-366) and pnCS-RPAP3 (133-255) plasmids. Cells were grown in LB medium containing 100 μg/ml of ampicillin, 25 μg/ml of chloramphenicol and 25 μg/ml of spectinomycin at 37°C under shaking. Protein expression was then induced with 0.5 mM IPTG for 16 h at 20°C once bacterial culture absorbance was of 0.6-0.8 at 600 nm (A_600_). Then, the cells were harvested by centrifugation for 30 min at 4,000 g at 4°C. The cell pellet was resuspended in 50 ml of purification buffer (25 mM HEPES pH 7.5, 300 mM NaCl, 0.5mM TCEP, 10 mM Imidazole) and sonicated. The complex was purified using TALON beads (Clontech) after nucleic acid precipitation using 0.05% of PolyEthylImine (PEI) and eluated by using PreScission (3C) Protease overnight on beads at 4°C to remove the N-terminal His_6_-tag. This step was followed by a preparative gel filtration (HiLoad 16/60 Superdex 75, GE Healthcare) on an AKTA prime system in 25 mM HEPES buffer (pH 7.5), 300 mM NaCl, 0.5 mM TCEP and 10 mM imidazole. Finally, the complex was concentrated to 11 mg/ml. For co-expression assays, protocol was the same, except we used gateway pnEA-3CH, pnCS and pnYK. Beads were directly resuspended in 2x Laemmli buffer and loaded on 15% SDS-PAGE instead of preparative gel filtration.

For the TECAN automated screen, *E. coli* BL21 (DE3) pRARE2 cells were co-transformed with the gateway pnEA-3CH and pnCS vectors and growth in Graffinity I buffer (2x LB, 0.5% Glucose). Expression was auto-induced by the addition of Graffinity II medium (v/v) (2x LB, 20 mM HEPES (pH7), 0.6% Lactose, 1 mM imidazole) overnight at 20°C, when absorbance reached 1.2 at 600 nm. Purification was performed with His_6_-tag Isolation Dynabeads (ThermoFischer Scientific) in low salt (LS: 20 mM Tris-HCl (pH8), 50 mM NaCl, 7 mM imidazole) or high salt (HS: 20 mM Tris-HCl (pH8), 500 mM NaCl, 7 mM imidazole) buffers. After three washes with LS or HS buffers, beads were resuspended in 2x Laemmli buffer and loaded on 15% SDS-PAGE.

### GST pull-downs

Total cellular extracts in RSB 100 buffer (100 mM NaCl, 10 mM Tris-HCl pH 7.5, 2.5 mM MgCl_2_, 0.01% NP40) were pre-cleared on Glutathione Sepharose beads for 2 h at 4°C. 4 µg of GST or of the GST-tagged protein of interest attached on Sepharose beads were incubated with 500 µl of pre-cleared cell extract for 2 h at 4°C on a rotating wheel. Beads were washed 5 times in RSB 200 buffer (200 mM NaCl, 10 mM Tris-HCl pH 7.5, 2.5 mM MgCl_2_, 0.05% NP40). After the last wash, beads were resuspended in SDS-PAGE loading dye and directly submitted to electrophoresis prior western-blotting.

### PLA and image acquisition

*In situ* proximity ligation assay (PLA) was performed as recommended by the manufacturer (DuolinkII kit, Olink Bioscience AB). Briefly, HeLa cells grown on coverslips were fixed in 1x PBS, 3% paraformaldehyde during 20 min and permeabilized for 5 min in a 1x PBS, 0.1% Triton X-100 solution. Primary antibodies were diluted in 1x antibody dilution buffer and incubated for 1 h at room temperature. The negative controls used only one of each primary antibody. Cells were washed three times for 5 min in 1x PBS. The PLA probes (Rabbit-MINUS and Mouse-PLUS) were incubated in a pre-heated humidity chamber for 1 h at 37°C. Subsequent steps were performed using the detection reagents green according to DuolinkII kit protocol. Finally, cells were incubated for 20 min with Alexa Fluor™ 546 Phalloidin (Thermo Fischer scientific) to detect the cytoskeleton. The Duolink mounting medium was supplemented with 10 μM TO-PRO-3 final to counterstain nuclei. Laser confocal microscopy was performed with a SP5-AOBS X Leica confocal microscope. Images from each channel were recorded separately and then merged with the ImageJ software.

### Co-immunoprecipitation and Western Blot

Cells are transfected in 10 cm plates with 15 µg of pcDNA3.1/nV5-DEST (Invitrogen) fused with TRBP and 30 µl of JetPEI (Polyplus transfection). After 48 h, cells were lyzed in 500 µl of HNTG buffer containing protease inhibitor cocktail (Roche) and incubated for 20 min at 4°C. Alternatively, T-Rex stable cell lines were treated with doxycycline to induce expression of the Flag-TRBP or Flag-RPAP3 proteins. Cellular debris were removed by centrifugation for 10 min at 9,000 *g*. Extracts were incubated on G-beads (ThermoFisher Scientific) coupled to 10 µg of anti-V5 or anti-Flag antibodies for 2 h at 4°C. For control IP, beads without antibodies were used. If necessary, extract was incubated with 15 µg of RNAse A (ThermoFisher Scientific). Beads were then washed three times with ice-cold HNTG before being resuspended in 2x Laemmli buffer. Inputs and pellets were loaded on 12% SDS-PAGE and transferred to activated-ethanol PVDF membrane (Protean Amersham). Membranes were blocked with 5% non fat dry milk in PBST (0.1% Tween-20 in PBS) and incubated with V5 or anti-RPAP3 primary antibody diluted in 5% non fat dry milk followed by incubation with secondary antibody conjugated to HRP. Enzymatic activity was detected using the ECL prime kit (Amersham).

### Luciferase assays

HEK-293T cells were grown on 96-well plates and co-transfected with 95 ng of plasmid expressing a HA-Tag Renilla luciferase (RL) in fusion with the protein of interest, and 5 ng of plasmid coding for the Firefly luciferase alone (FL) with 0.3 μl of JetPrime (Ozyme). After 48 h, cells were extracted in 50 µl of ice-cold 1x HNTG buffer (20 mM HEPES, pH 7.9, 150 mM NaCl, 1% Triton X-100, 10% glycerol, 1 mM MgCl_2_, 1 mM EGTA) containing protease inhibitor cocktail (Roche) and incubated at 4 °C for 15 min. RL and FL activities were measured on 96-well plates using 2 µl of cell extract containing 8 µl of 1x PLB (Promega) and the dual-luciferase assay kit (Promega). Values obtained for RL were normalized to FL values. Experiments were done at least in triplicate. For geldanamycin (GA) experiments, drug was added 16 h before extraction to a final concentration of 2 µM.

### LUMIER IP

HEK-293T cells were grown in 24-well plates and co-transfected with 450 ng of the RL fusion and 50 ng of the 3x FLAG-FL fusion. After 48 h, cells were extracted in 500 μl of ice-cold HNTG containing protease inhibitor cocktail (Roche), incubated for 15 min at 4 °C and spun down at 4 °C at 20,000 *g* for 15 min. 100 µl of the extract were dispatched in two wells in a 96-well plate, with one well being coated with anti-FLAG antibody (10 μg/ml in 1x PBS, F1804 Sigma-Aldrich), and one control well without antibodies. Plates were incubated for 3 h at 4 °C, and then washed 5 times with 300 μl of ice-cold HNTG, for 10 min at 4 °C for each wash. After the last wash, 10 μl of 1x PLB (Promega) was added in each well. To measure the input, 2 μl of extract and 8 μl of 1x PLB were mixed in new wells. Plates were then incubated for 5 min at room temperature, and FL and RL luciferase activities were measured in IP and input wells, using the dual luciferase kit (Promega). Experiments were done at least in triplicate. Co-IP efficiency was defined as the RL/FL ratio in the pellet, divided by the RL/FL ratio in the input. Unless stated otherwise, statistical significance was evaluated using Z-test assaying whether the co-IP efficiency in the anti-FLAG IP was more than 11 times higher than the mean values obtained in the control IP, done without antibodies.

### Nuclear Magnetic Resonance

A perdeuterated ^13^C/^15^N labeled sample of the complex between RPAP3-TPR1 (i.e. fragment 133-155 of human RPAP3) and TRBP-dsRBD3 (i.e. fragment 262-366 of human TRBP) was prepared as the X-ray sample, except that bacteria were initially grown in a minimal M9 medium supplemented with ^13^C-D6-glucose, ^15^N ammonium chloride and 50% D_2_O. The final sample was concentrated at 1 mM in 10 mM NaPi, 150 mM NaCl, 5 mM ^2^H DTT and is stable for about 2 days. ^1^H-^15^N HSQC, HNCO, HNCA, HNCACB, CBCACONH and a ^1^H-^15^N NOESY-HSQC (with a mixing time of 120 ms) spectra were recorded at 303 K on 600 and 950 MHz spectrometers equipped with cryoprobes. Assignment of backbone resonances was performed with CARA (61). The chemical shift data were derived into secondary structures using TALOS+ (62). Free and TRBP-dsRBD3 bound states of RPAP3-TRP1 were compared using a previous assignment performed in the same experimental conditions (BMRB entry19758,(39) and a composite ^1^H-^15^N chemical shift calculated for each residue as follow: 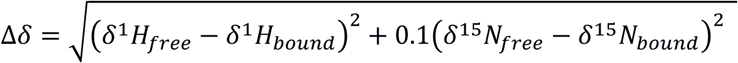.The assignment of the free RPAP3-TPR1 was obtained using a protein fragment holding four non-native residues located at its N-terminal position. To avoid a bias on chemical shift perturbations generated by this difference in the primary structure, residues in proximity of the N-terminal tail of RPAP3-TPR1 were discarded from the analysis. The residues in RPAP3-TPR1 with a Δδ value superior to the centile 80 value were considered as significantly perturbed upon binding of TRBP-dsRBD3.

### Protein crystallization, X-Ray data collection and structure determination

#### Crystallization and X-ray data collection

Crystals of the complex between RPAP3 (residues 133-255) and TRBP (residues 262-366) were grown by vapor diffusion in hanging drops. Drops were made at 293 K by mixing 2 µl of the protein solution at 11 mg/ml and 2 µl of a reservoir solution containing 18% (w/v) PEG 3,350 and 8% (v/v) Tacsimate™ at pH 6.0. Crystals belong to space group *P*2_1_2_1_2 with unit-cell parameters a = 39.8 Å, b = 158.1 Å and c = 32.7 Å. Assuming one heterodimer in the asymmetric unit, the packing density *V*_M_ is 2.00 Å^3^.Da^-1^ and the solvent content is 38.4%. Crystals were flash frozen in liquid nitrogen in the mother liquor with addition of 25% glycerol as cryoprotectant. A native data set at 1.49 Å resolution was collected at 100 K on beamline ID29 at the European Synchrotron Radiation Facility (ESRF, Grenoble), with incident radiation at a wavelength of 1.033 Å and a crystal-to-detector distance of 207 mm. Diffraction spots were recorded on a Pilatus 6M-F detector with a 0.1° oscillation and a 0.04 second exposure per image. Data were indexed and scaled using XDS (63) and indexed intensities were converted to structure factors using TRUNCATE in the CCP4 suite (64) without any σ cut-off.

#### Crystal structure determination

The crystal structure of the RPAP3:TRBP complex was solved by molecular replacement with the program PHASER (65) using the coordinates of RPAP3 from the crystal structure of RPAP3 bound to a HSP90 peptide ((35); PDB 4CGV) and the coordinates of TRBP bound to Dicer ((10); PDB 4WYQ). A single solution was obtained with LLG = 795 and TFZ = 18.2. Building of the model was performed using Coot (66), and the refinement of the crystal structure was performed in the range 40-1.49 Å using REFMAC5 (67). A total of 4% of the native data were selected for *R*_free_ calculations. The model was refined to the final *R*_factor_ of 17.8% and *R*_free_ of 21.1% **(Table 1)** and includes residues 133-249 of RPAP3, residues 263-365 of TRBP and 243 water molecules. Because of the lack of density, residues 250-265 of RPAP3 and residues 262 and 366 of TRBP were not built. They were probably too flexible in the complex to generate a clear electron density. Coordinates of the RPAP3:TRBP structure have been deposited in the Protein Data Bank (accession number 6ZBK). Over 97% of the residues were within the most favored regions, and no residue was within the disallowed regions in a Ramachandran plot, as defined by PROCHECK (68). Averaged B factors were of 15.3 Å^2^ for the protein atoms, 26.5 Å^2^ for water molecules, and 16.7 Å^2^ for the whole structure. Figures were prepared using PyMOL (The PyMOL Molecular Graphics System, Version 1.8 Schrödinger, LLC).

## Supporting information

Supplementary results, figures and tables

## ACCESSION CODE

The TRBP-dsRBD 3:RPAP3-TPR1 complex coordinates and structure factors have been deposited in the PDB with accession code 6ZBK.

*Note: Supplementary information is available in the online version of the paper*.

## ACKNOWLEDGMENTS

Centre National de la Recherche Scientifique; Université de Lorraine; Y.A. was a fellow from the French Ministry of Education and Research; Research of M.R. was supported by the Université de Lorraine and the CNRS (‘Chaire d’Excellence’ program), by the Pôle BMS of the Université de Lorraine (grant CS-UL 2018 AAP-BMS_003_162_INCITATIF_IMoPA_Rederstorff M.) and by the Ligue contre le Cancer (Comités 54, 57). Financial support from the IR-RMN-THC Fr3050 CNRS for conducting the research is gratefully acknowledged.

We are thankful to Sébastien Pfeffer, Natacha Dreumont and members of the IMoPA team 1 for helpful discussions.

We thank the platforms “Biophysique et Biologie Structurale” (B2S) and “Imagerie et de Biophysique Cellulaire” (PTIBC) of UMS 2008 IBSLor for access to X-ray crystallography, NMR facilities and confocal microscopes. We are grateful to the synchrotron ESRF (Grenoble, France) for access to the ID29 beamline. Séverine Massenet is acknowledged for the PIH1D1 and RUVBL1/2 antibodies.

## AUTHOR CONTRIBUTIONS

C.B., M.R. initiated the project. Y.A., B.C., M.R. designed the experiments with inputs from all authors. Y.A.,, V.B-I., M.Q., M-E.C., M.R. performed the experiments. C.C. performed the crystallogenesis the X-ray structural analysis. Y.A., M.Q., performed the NMR experiments. All authors contributed to data interpretation. Y.A., C.C., E.B., B.C., M.R. wrote the manuscript with inputs from M.Q., C.V., C.B., X.M.. All authors proofread the manuscript.

## FIGURE LEGENDS

**Table 1.** Crystallographic data collection and refinement statistics.

