## Supplementary results, figures and tables for "Structure of the RPAP3:TRBP interaction reveals an involvement of the HSP90/R2TP chaperone complex in dsRNA pathways"

**SUPPLEMENTARY INFORMATION**

*The 3D structure of RPAP3 in the RPAP3:TRBP complex displays a TPR fold*

The crystal structure of the complex between RPAP3 (residues 133-249) and TRBP (residues 263-365) was refined to the final $R_{factor}$ of 17.8% and $R_{free}$ of 21.1% at 1.49 Å resolution. The asymmetric unit contains one RPAP3:TRBP heterodimer and 243 water molecules.

Superimposition of the crystal structure of RPAP3 (residues 133-249) bound to TRBP with the crystal structure of RPAP3 bound to the C-terminal tail peptide (SRMEEVD) of HSP90 (35) yields a root-mean-square deviation (RMSD) of C$\alpha$ positions of 0.59 Å **(Supp. Fig. 5 a)**.

*The 3D structure of TRBP in the RPAP3:TRBP complex contains a typical dsRBD fold*

The dsRBD core of TRBP bound to RPAP3 is structurally similar to the dsRBD domain of TRBP in complex with Dicer, with a backbone RMSD of 0.78 Å over 74 residues **(Supp. Fig. 5 b)**. The result of this comparison suggests that RPAP3 may not induce specific significant conformational changes on TRBP upon binding.

However, the 3D structure of the free TRBP is still not available and a comparison between the structure of TRBP in a free state and the structure of TRBP bound to RPAP3 would be necessary to confirm this hypothesis. Interestingly, an N-terminal extension (residues 263-288) beyond the canonical dsRBD domain of TRBP is observed in the crystal structure of the RPAP3:TRBP complex and consists of 3 helices H1, H2 and H3 **(Fig. 4, Supp. Fig. 5 b, c)**. This N-terminal extension was shown to be partially disordered in the crystal structure of TRBP bound to protein Dicer (10) since only $\alpha$-helix H1 was partially modeled as a separate chain of alanine. In the RPAP3:TRBP structure, the $\alpha$-helix H1 is located at the opposite side of the β-sheet and lies at the surface of both $\alpha$-helices H4 and H5. The axis of $\alpha$-helix H2 is perpendicular to the axis of $\alpha$-helix H1, which allows $\alpha$-helix H2 to position in a cleft that is delineated by the C-terminus of $\alpha$-helices H4 and H5 as well as loops H4-β1 and β2-β3. The hydrophobic residues W265 and L268 from $\alpha$-helix H1 as well as I276 and L279 from $\alpha$-helix H2 contribute to a large hydrophobic cluster of TRBP **(Supp. Fig. 9 f)** that includes residues L356, Y358, L359 and M362 from $\alpha$-helix H5 as well as V297 and L301 from $\alpha$-helix H4, and F307 in the loop H4-β1.

*RPAP3 interacts via its first TPR domain to the dsRBD3 domain of TRBP*

The RPAP3:TRBP interaction buries 1,517 Å$^2$ of molecular surface, and it involves 7% of the total solvent-accessible surface area of the first TPR domain of RPAP3 and 9% of the total solvent-accessible surface area of the TRBP core. The dsRBD3 domain of TRBP binds to the central part of the TPR1 domain of RPAP3 **(Fig. 4a)** that includes α-helices α2, α3, and α4 while no interaction is made with the N-terminal α-helix α1 as well as the three C-terminal α-helices α5, α6, and α7 of the TPR domain of RPAP3. On the other hand, the RPAP3:TRBP interaction buries the C-terminal region of the dsRBD domain of TRBP and the interface involves mainly the C-terminal α-helix H5 as well as the β-strands β2 and β3 of TRBP. As shown in **Fig. 4 a**, the C-terminal α-helix H5 of TRBP lies in a cleft that is delineated by α-helices α2, α3, and α4 of RPAP3. Interestingly, both α-helices H1 and H2 that are located in the N-terminal extension (residues 263-288) beyond the canonical dsRBD domain of TRBP are not involved in the interface of the heterodimer. Otherwise, only one residue (L284) from this N-terminal extension, that is located in the loop H2- H3, is close to RPAP3.

*Hydrophobic contacts, salt-bridges and hydrogen bonds at the RPAP3:TRBP interface*

Several hydrogen-bonds as well as some hydrophobic contacts and ionic interactions are observed between side-chain of residues involved at the interface of the RPAP3:TRBP heterodimer. First, residue D150 from RPAP3 is hydrogen-bonded both with S318 and S320 from TRBP **(Supp. Fig. 8 c)**. This polar interaction connects the N-terminal part of α-helix α2 of RPAP3 to the loop β1-β2 of TRBP. Second, three hydrophobic residues from RPAP3 form hydrophobic contacts with 4 hydrophobic residues of TRBP. Indeed, residues M160 and L192 from RPAP3 form a small hydrophobic cluster with residues Y358 and I361 of TRBP **(Sup. Fig. 8 a)**. This cluster allows the interaction of the C-terminal α- helix H5 of TRBP with the N-terminal part of both α-helices α2 and α4 of RPAP3. Another hydrophobic contact involves residue V185 of RPAP3 and residues L326 and V336 of TRBP **(Supp. Fig. 8 b)**. This intermolecular interaction connects the N-terminal part of α-helix α4 of RPAP3 to both β-strands β2 and β3 of TRBP.

This hydrophobic cluster is essential for the RPAP3:TRBP interaction since the mutation of V185A in RPAP3 leads to the loss of interaction of RPAP3 with TRBP. In addition, the backbone nitrogen of

V336 of TRBP forms a hydrogen bond with the side-chain of S188 of RPAP3. Finally, a small cluster of electronic interactions is observed at the heterodimer interface **(Fig. 6 a)**. It involves the acidic residues D154 and D161 from RPAP3 and the positively-charged residues R353 and R354 of TRBP. Moreover, residue D161 from RPAP3 also forms a hydrogen-bond with residue Q357 of TRBP while residue R354 from TRBP is also hydrogen-bonded to residue T157 of RPAP3. This network of polar interactions allows the interaction of α-helix α2 of RPAP3 with the C-terminal α-helix H5 of TRBP. Interestingly, the α-helix α2 of RPAP3 interacts via hydrophobic contacts as well as ionic interactions and hydrogen bonds with the C-terminal α-helix H5 of TRBP and seems to be a crucial secondary-structure element for the RPAP3:TRBP complex formation. The mutation of D161A or T157A in RPAP3 leads to the disruption of the RPAP3:TRBP complex. Likely, the mutation of R354 in TRBP destabilizes the RPAP3:TRBP interaction. Such results from mutation experiments confirm the importance to form a polar cluster at the interface between RPAP3 and TRBP.

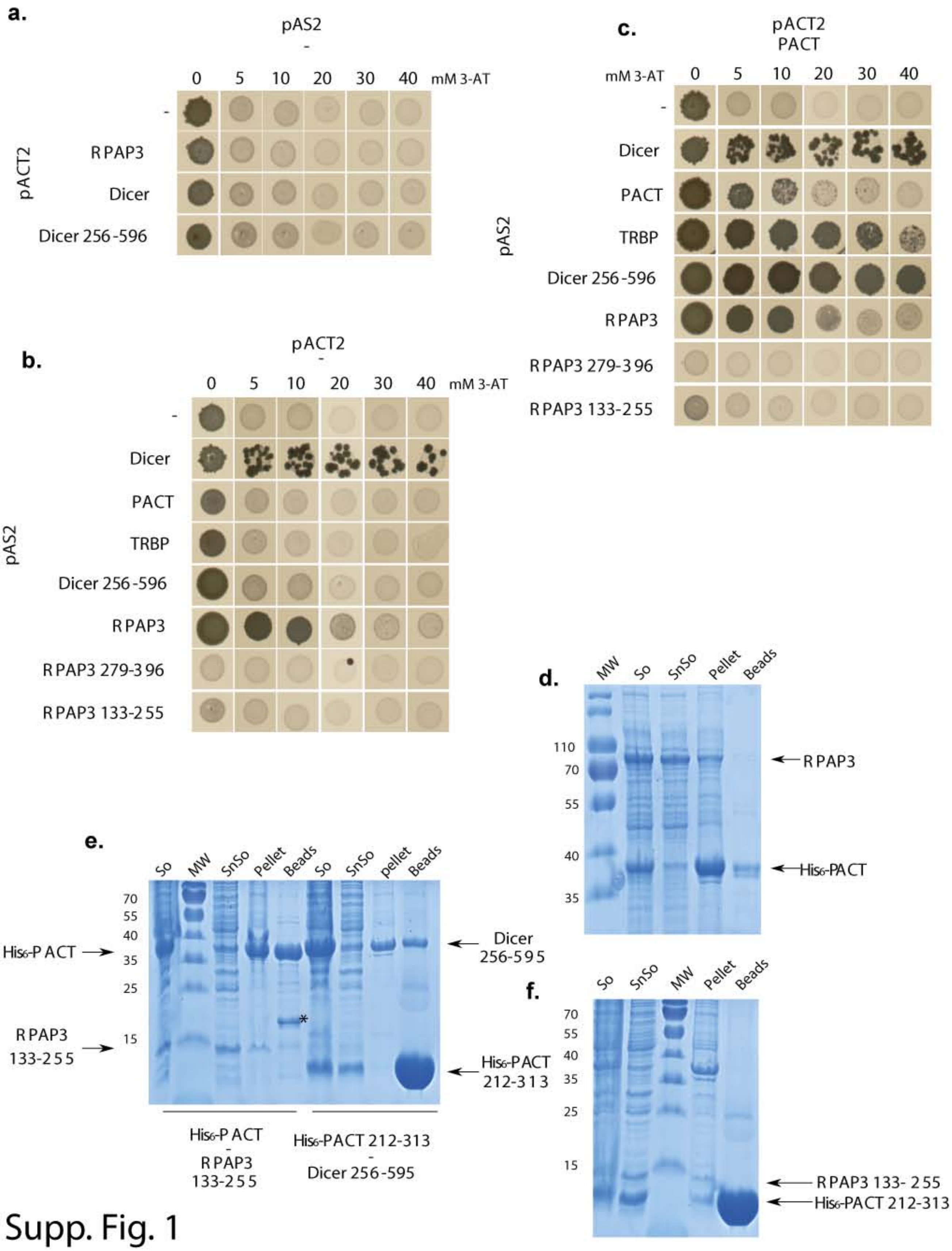
a.
pAS2
-
0 5 10 20 30 40 mM 3-AT
pACT2
-
RPAP3
Dicer
Dicer 256-596
b.
pACT2
-
0 5 10 20 30 40 mM 3-AT
pAS2
-
Dicer
PACT
TRBP
Dicer 256-596
RPAP3
RPAP3 279-396
RPAP3 133-255
c.
pACT2
PACT
mM 3-AT 0 5 10 20 30 40
pAS2
-
Dicer
PACT
TRBP
Dicer 256-596
RPAP3
RPAP3 279-396
RPAP3 133-255
d.
MW So SnSo Pellet Beads
110
70
55
40
35
RPAP3
His6-PACT
e.
So MW SnSo Pellet Beads So SnSo pellet Beads
70
55
40
35
25
15
His6-PACT
RPAP3 133-255
Dicer 256-595
His6-PACT 212-313
*
His6-PACT - RPAP3 133-255
His6-PACT 212-313 - Dicer 256-595
f.
So SnSo MW Pellet Beads
70
55
40
35
25
15
RPAP3 133- 255
His6-PACT 212-313

Supp. Fig. 1

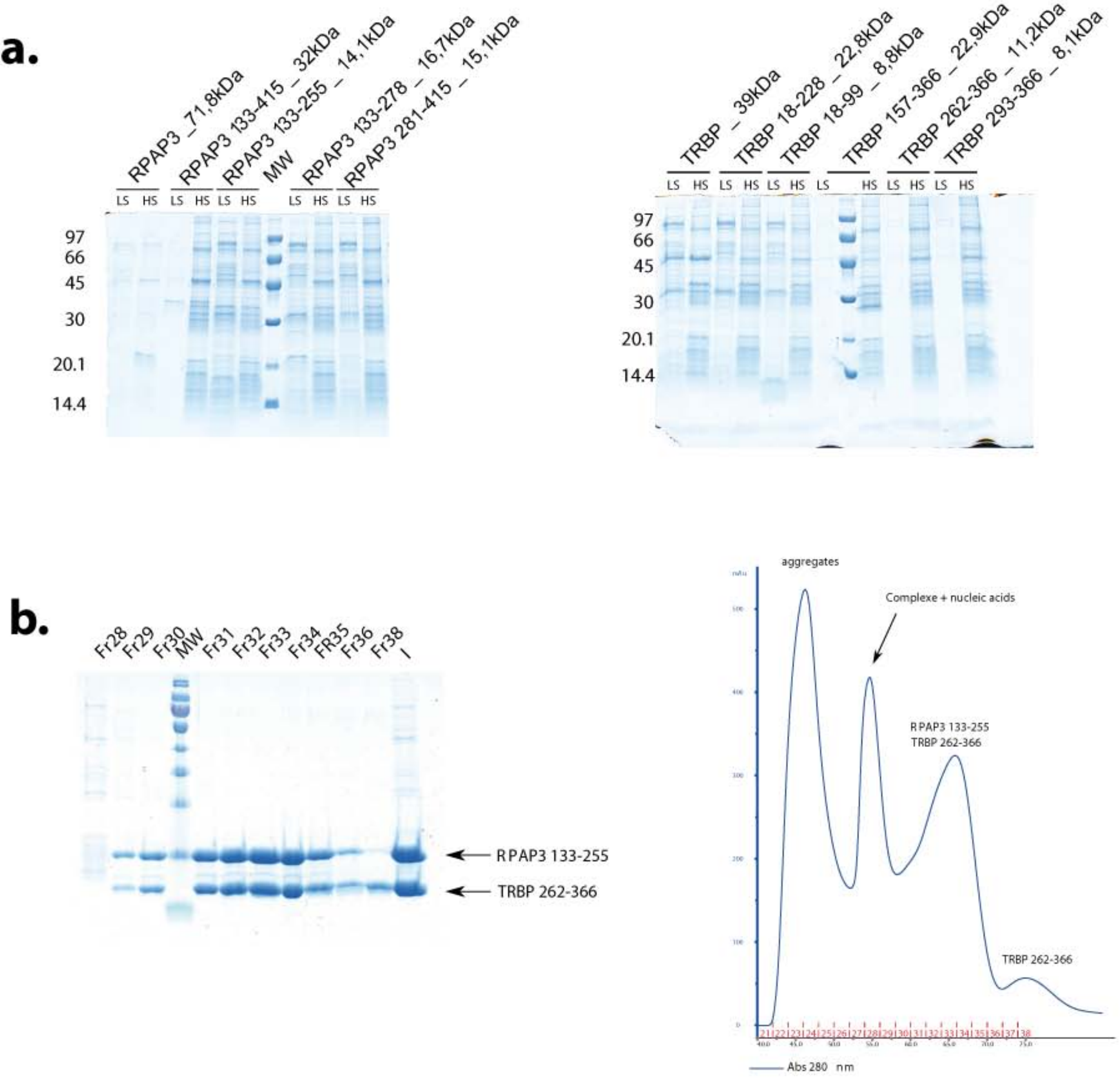
a.
RPAP3 _71,8kDa
RPAP3 133-415 _ 32kDa
RPAP3 133-255 _ 14,1kDa
MW
RPAP3 133-278 _ 16,7kDa
RPAP3 281-415 _ 15,1kDa
LS
HS
97
66
45
30
20.1
14.4
TRBP _ 39kDa
TRBP 18-228 _ 22,8kDa
TRBP 18-99 _ 8,8kDa
TRBP 157-366 _ 22,9kDa
TRBP 262-366 _ 11,2kDa
TRBP 293-366 _ 8,1kDa
b.
Fr28
Fr29
Fr30
MW
Fr31
Fr32
Fr33
Fr34
FR35
Fr36
Fr38
I
RPAP3 133-255
TRBP 262-366
aggregates
Complexe + nucleic acids
RPAP3 133-255
TRBP 262-366
TRBP 262-366
Abs 280 nm

Supp. Fig. 2

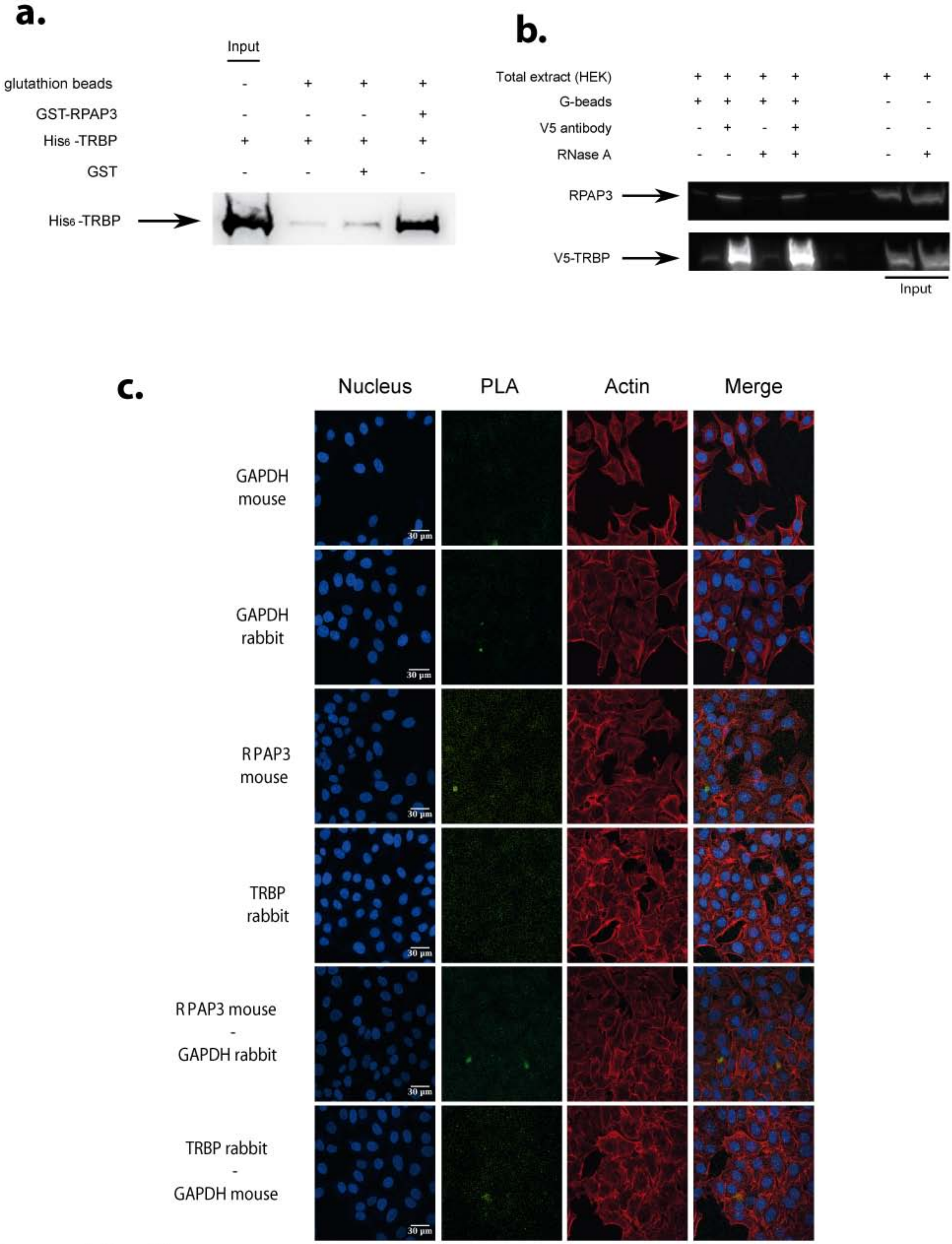
a.
Input
glutathion beads
GST-RPAP3
His6 -TRBP
GST
His6 -TRBP
b.
Total extract (HEK)
G-beads
V5 antibody
RNase A
RPAP3
V5-TRBP
Input
c.
Nucleus
PLA
Actin
Merge
GAPDH mouse
GAPDH rabbit
RPAP3 mouse
TRBP rabbit
RPAP3 mouse - GAPDH rabbit
TRBP rabbit - GAPDH mouse
30 µm

Supp. Fig. 3

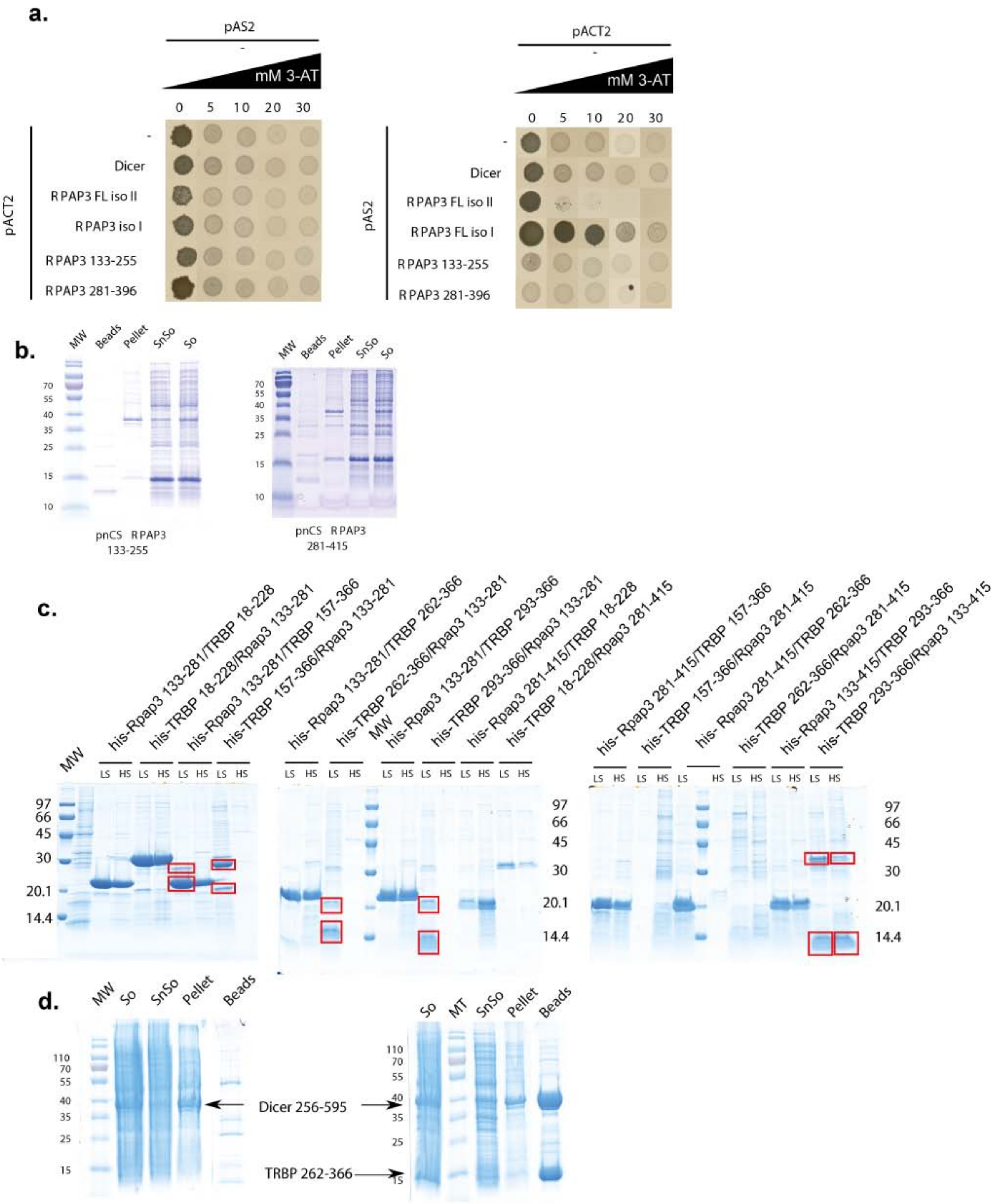
a.
pAS2
pACT2
mM 3-AT
0 5 10 20 30
Dicer
RPAP3 FL iso II
RPAP3 iso I
RPAP3 FL iso I
RPAP3 133-255
RPAP3 281-396
b.
MW Beads Pellet SnSo So
pnCS RPAP3 133-255
pnCS RPAP3 281-415
c.
his-Rpap3 133-281/TRBP 18-228
his-TRBP 18-228/Rpap3 133-281
his-Rpap3 133-281/TRBP 157-366
his-TRBP 157-366/Rpap3 133-281
his-Rpap3 133-281/TRBP 262-366
his-TRBP 262-366/Rpap3 133-281
his-Rpap3 133-281/TRBP 293-366
his-TRBP 293-366/Rpap3 133-281
his-Rpap3 281-415/TRBP 18-228
his-TRBP 18-228/Rpap3 281-415
his-Rpap3 281-415/TRBP 157-366
his-TRBP 157-366/Rpap3 281-415
his-Rpap3 281-415/TRBP 262-366
his-TRBP 262-366/Rpap3 281-415
his-Rpap3 133-415/TRBP 293-366
his-TRBP 293-366/Rpap3 133-415
LS HS
97 66 45 30 20.1 14.4
d.
MW So SnSo Pellet Beads
So MT SnSo Pellet Beads
Dicer 256-595
TRBP 262-366

Supp. Fig. 4

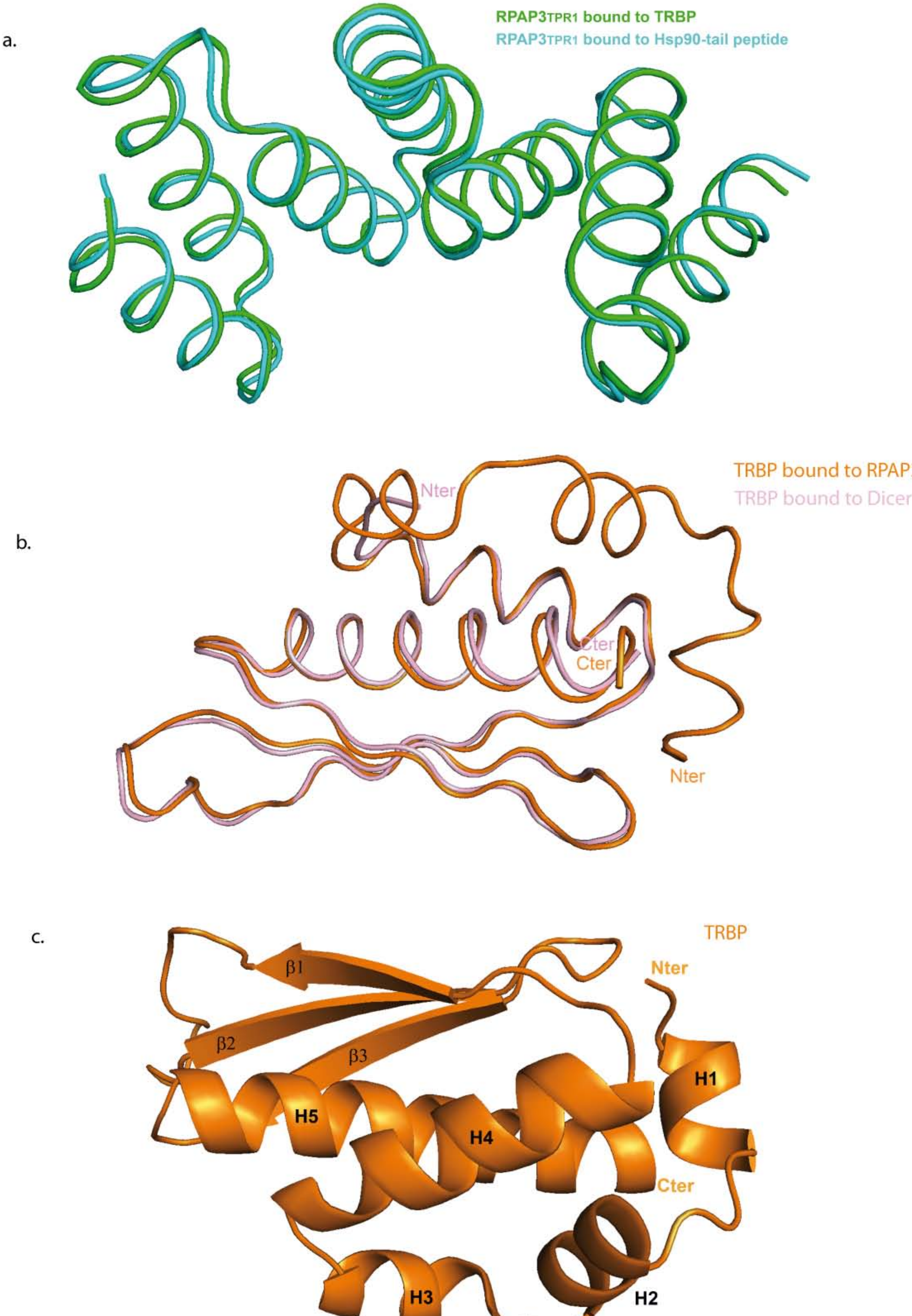
a.
RPAP3TPR1 bound to TRBP
RPAP3TPR1 bound to Hsp90-tail peptide
b.
TRBP bound to RPAP3
TRBP bound to Dicer
Nter
Cter
Cter
Nter
c.
TRBP
Nter
β1
β2
β3
H1
H5
H4
Cter
H3
H2

Supp. Fig. 5

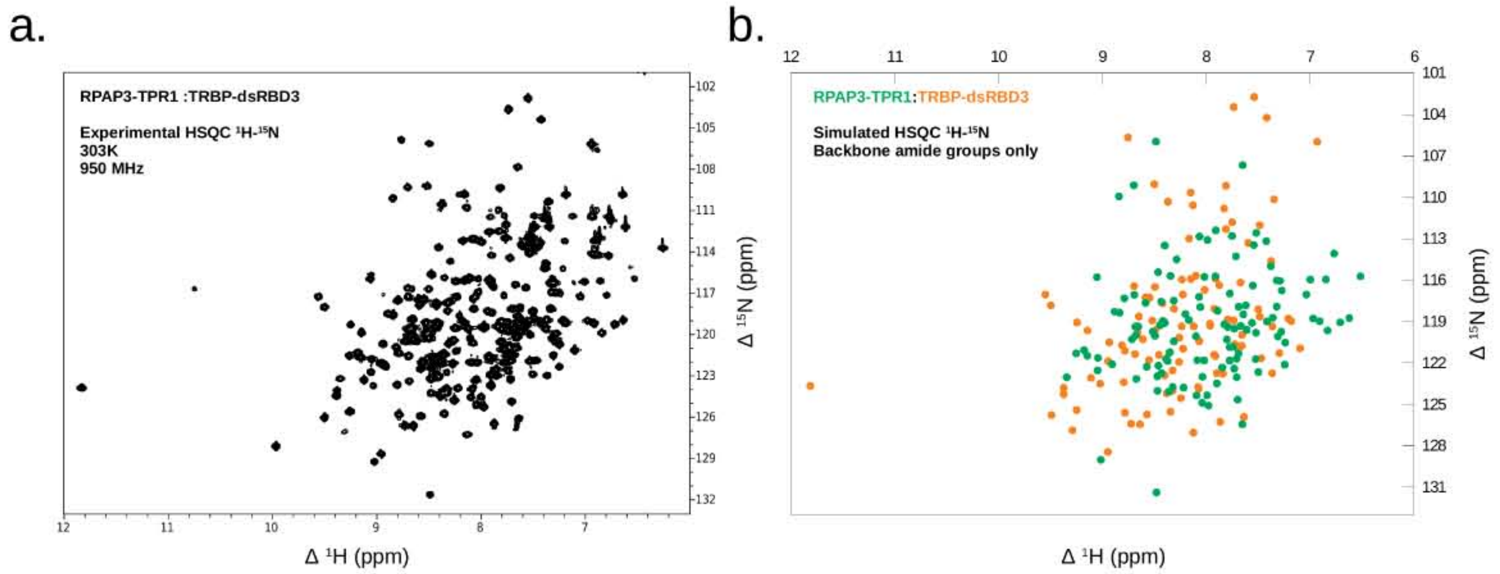

c.

RPAP3-TPR1 (fragment 133-255)

| Residue | ¹H (ppm) | ¹⁵N (ppm) | Residue | ¹H (ppm) | ¹⁵N (ppm) |
|---|---|---|---|---|---|
| V135 | 8.372 | 121.857 | N197 | 7.914 | 115.722 |
| L136 | 8.324 | 123.737 | R198 | 8.674 | 123.149 |
| K137 | 8.400 | 119.102 | S199 | 8.788 | 117.315 |
| E138 | 7.689 | 117.921 | Y200 | 7.500 | 122.658 |
| K139 | 8.386 | 121.716 | T201 | 8.040 | 124.884 |
| G140 | 8.485 | 105.961 | K202 | 7.552 | 116.396 |
| N141 | 8.494 | 120.031 | A203 | 7.361 | 118.712 |
| K142 | 8.037 | 123.003 | Y204 | 7.000 | 115.954 |
| Y143 | 7.891 | 118.220 | S205 | 7.716 | 114.267 |
| F144 | 9.044 | 121.628 | R206 | 7.670 | 119.311 |
| K145 | 8.312 | 120.428 | R207 | 8.361 | 124.085 |
| Q146 | 7.380 | 114.989 | G208 | 8.697 | 109.121 |
| G147 | 7.650 | 107.672 | A209 | 7.698 | 124.659 |
| K148 | 7.732 | 122.414 | A210 | 8.471 | 122.969 |
| Y149 | 6.625 | 118.741 | R211 | 9.053 | 115.779 |
| D150 | 8.679 | 120.001 | F212 | 9.048 | 122.542 |
| E151 | 9.344 | 123.032 | A213 | 7.708 | 123.033 |
| A152 | 8.021 | 121.818 | L214 | 7.273 | 116.753 |
| I153 | 8.434 | 117.559 | Q215 | 7.989 | 113.091 |
| D154 | 7.993 | 124.340 | K216 | 8.432 | 122.732 |
| C155 | 7.774 | 116.971 | L217 | 7.324 | 117.919 |
| Y156 | 8.883 | 118.279 | E218 | 8.835 | 118.360 |
| T157 | 8.345 | 115.688 | E219 | 8.655 | 119.362 |
| K158 | 8.087 | 121.801 | A220 | 8.461 | 122.185 |
| G159 | 8.841 | 109.925 | K221 | 8.313 | 117.485 |
| M160 | 8.094 | 124.351 | K222 | 7.324 | 116.030 |
| D161 | 6.771 | 114.068 | D223 | 7.526 | 121.721 |
| A162 | 6.909 | 118.972 | Y224 | 8.058 | 117.953 |
| D163 | 7.644 | 118.472 | E225 | 8.683 | 119.371 |
| Y165 | 7.909 | 112.393 | R226 | 7.847 | 122.325 |
| N166 | 6.972 | 118.781 | V227 | 7.827 | 119.197 |
| V168 | 7.709 | 121.689 | L228 | 7.292 | 116.047 |
| L169 | 6.517 | 115.721 | E229 | 7.561 | 119.098 |
| T171 | 6.850 | 115.941 | L230 | 7.417 | 118.989 |
| N172 | 8.219 | 123.788 | E231 | 9.146 | 121.468 |
| R173 | 8.470 | 124.017 | N233 | 8.401 | 113.486 |
| A174 | 8.907 | 122.091 | N234 | 7.611 | 119.602 |
| S175 | 8.286 | 114.494 | F235 | 8.480 | 131.373 |
| A176 | 7.977 | 125.092 | E236 | 8.350 | 121.215 |
| Y177 | 8.693 | 117.057 | A237 | 8.715 | 120.276 |
| F178 | 9.253 | 121.304 | T238 | 7.519 | 112.570 |
| R179 | 7.745 | 120.847 | N239 | 7.802 | 120.302 |
| L180 | 7.038 | 117.040 | E240 | 8.515 | 119.731 |
| K181 | 7.423 | 113.158 | L241 | 8.167 | 118.885 |
| K182 | 7.793 | 119.389 | R242 | 7.312 | 120.074 |
| F183 | 6.712 | 119.061 | K243 | 7.280 | 119.733 |
| A184 | 9.177 | 121.071 | I244 | 8.573 | 122.272 |
| V185 | 6.833 | 119.652 | S245 | 8.464 | 115.426 |
| A186 | 7.732 | 119.521 | Q246 | 7.775 | 120.846 |
| E187 | 8.585 | 117.651 | A247 | 7.898 | 123.482 |
| S188 | 7.544 | 113.455 | L248 | 8.446 | 119.276 |
| D189 | 9.016 | 129.010 | A249 | 7.690 | 121.304 |
| C190 | 8.069 | 117.209 | S250 | 7.754 | 112.798 |
| N191 | 8.420 | 118.949 | K251 | 7.776 | 121.828 |
| L192 | 7.246 | 122.121 | E252 | 8.100 | 120.063 |
| A193 | 7.523 | 119.812 | N253 | 8.202 | 118.452 |
| V194 | 8.066 | 112.835 | S254 | 8.018 | 115.752 |
| A195 | 7.240 | 120.519 | Y255 | 7.655 | 126.431 |
| L196 | 7.621 | 117.824 | | | |

TRBP-dsRBD3 (fragment 262-366)

| Residue | ¹H (ppm) | ¹⁵N (ppm) | Residue | ¹H (ppm) | ¹⁵N (ppm) |
|---|---|---|---|---|---|
| H260 | 8.513 | 119.768 | D313 | 8.640 | 126.441 |
| M261 | 8.325 | 122.542 | I314 | 7.638 | 125.902 |
| G262 | 8.151 | 109.642 | E315 | 8.949 | 128.435 |
| C263 | 8.360 | 121.305 | E316 | 7.966 | 119.156 |
| T264 | 8.421 | 117.688 | L317 | 8.724 | 126.405 |
| W265 | 8.260 | 120.161 | S318 | 9.549 | 117.043 |
| D266 | 8.235 | 115.954 | L319 | 9.373 | 123.800 |
| S267 | 8.016 | 116.711 | S320 | 7.828 | 110.803 |
| L268 | 7.841 | 122.779 | G321 | 8.129 | 110.575 |
| R269 | 8.149 | 115.904 | L322 | 7.793 | 118.848 |
| N270 | 7.598 | 113.297 | C323 | 8.122 | 119.352 |
| S271 | 7.376 | 114.619 | Q324 | 9.497 | 117.823 |
| V272 | 8.490 | 119.631 | C325 | 8.686 | 121.358 |
| G273 | 7.811 | 109.157 | L326 | 8.783 | 125.593 |
| E274 | 8.262 | 121.925 | V327 | 8.946 | 121.875 |
| K275 | 11.811 | 123.654 | E328 | 8.571 | 125.727 |
| I276 | 9.491 | 125.781 | L329 | 9.374 | 124.255 |
| L277 | 8.381 | 124.186 | S330 | 8.811 | 120.717 |
| S278 | 7.911 | 115.884 | T331 | 7.348 | 110.153 |
| L279 | 7.901 | 122.741 | Q332 | 7.737 | 118.918 |
| R280 | 8.546 | 117.272 | A334 | 8.401 | 122.886 |
| S281 | 7.752 | 111.803 | T335 | 7.659 | 120.759 |
| C282 | 7.364 | 119.361 | V336 | 8.125 | 127.039 |
| S283 | 8.696 | 116.436 | C337 | 8.783 | 121.110 |
| L284 | 9.290 | 126.879 | H338 | 9.023 | 123.501 |
| G285 | 8.753 | 105.667 | S340 | 7.812 | 112.300 |
| S286 | 7.922 | 118.077 | A341 | 8.935 | 120.502 |
| L287 | 7.471 | 121.651 | T342 | 8.502 | 109.052 |
| G288 | 7.736 | 103.465 | T343 | 6.934 | 105.979 |
| A289 | 7.297 | 121.296 | R344 | 9.110 | 123.074 |
| L290 | 7.210 | 118.779 | E345 | 9.144 | 119.650 |
| G291 | 8.369 | 110.321 | A346 | 7.710 | 120.837 |
| A293 | 7.725 | 120.642 | A347 | 8.245 | 124.538 |
| C294 | 8.585 | 117.246 | R348 | 7.500 | 118.117 |
| C295 | 7.876 | 116.365 | G349 | 7.419 | 104.229 |
| R296 | 7.366 | 122.741 | E350 | 7.925 | 121.404 |
| V297 | 7.657 | 119.965 | A351 | 8.079 | 123.738 |
| L298 | 8.237 | 119.349 | A352 | 8.407 | 118.997 |
| S299 | 8.101 | 115.672 | R353 | 8.611 | 120.276 |
| E300 | 8.457 | 121.416 | R354 | 7.963 | 119.293 |
| L301 | 7.914 | 121.497 | A355 | 8.345 | 125.533 |
| S302 | 8.226 | 117.033 | L356 | 8.649 | 118.633 |
| E303 | 7.098 | 120.936 | Q357 | 8.072 | 123.835 |
| E304 | 7.182 | 118.974 | Y358 | 8.067 | 120.423 |
| Q305 | 8.165 | 112.982 | L359 | 8.495 | 116.494 |
| A306 | 7.486 | 118.647 | K360 | 8.520 | 118.924 |
| F307 | 7.486 | 112.017 | I361 | 8.224 | 120.962 |
| H308 | 9.245 | 119.066 | M362 | 8.311 | 118.079 |
| V309 | 8.323 | 123.990 | A363 | 8.661 | 119.779 |
| S310 | 8.552 | 121.786 | G364 | 7.540 | 102.722 |
| Y311 | 8.796 | 123.398 | S365 | 7.672 | 116.173 |
| L312 | 9.249 | 125.404 | K366 | 7.867 | 126.266 |

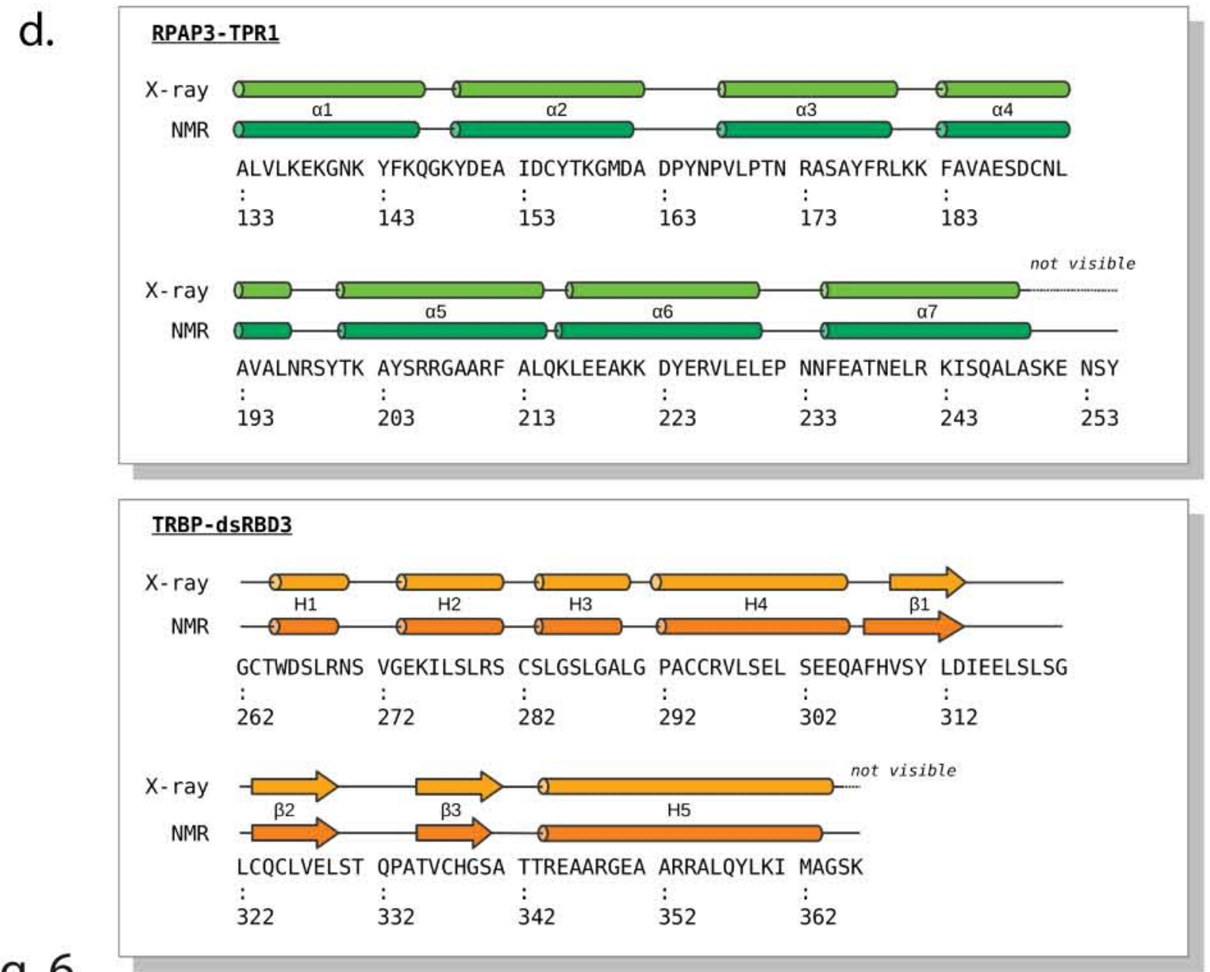

Supp. Fig. 6

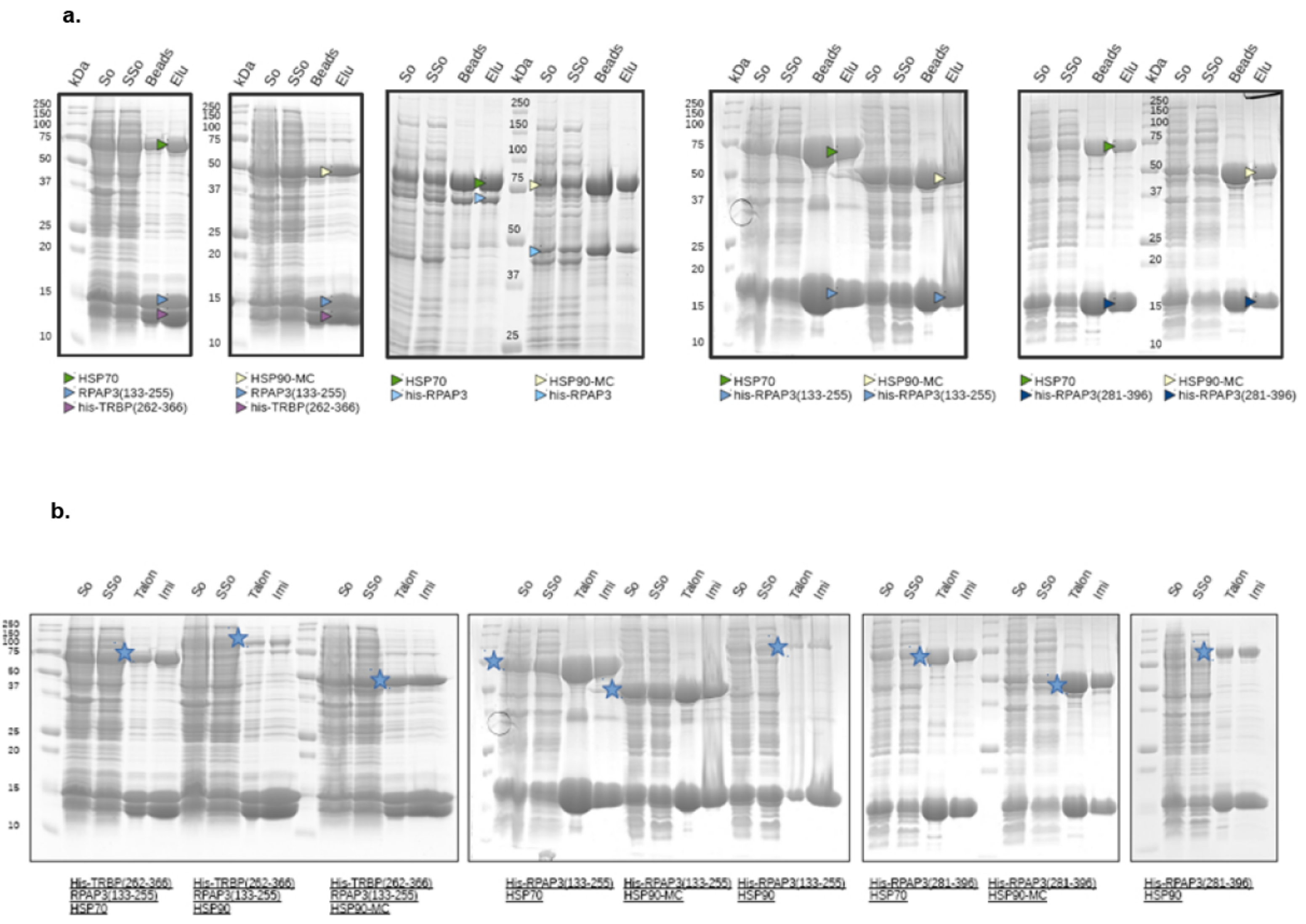
a.
kDa
So
SSo
Beads
Elu
HSP70
RPAP3(133-255)
his-TRBP(262-366)
HSP90-MC
his-RPAP3
his-RPAP3(133-255)
his-RPAP3(281-396)
b.
Talon
Imi
His-TRBP(262-366)
RPAP3(133-255)
HSP70
HSP90
HSP90-MC
His-RPAP3(133-255)
His-RPAP3(281-396)

Supp. Fig. 7

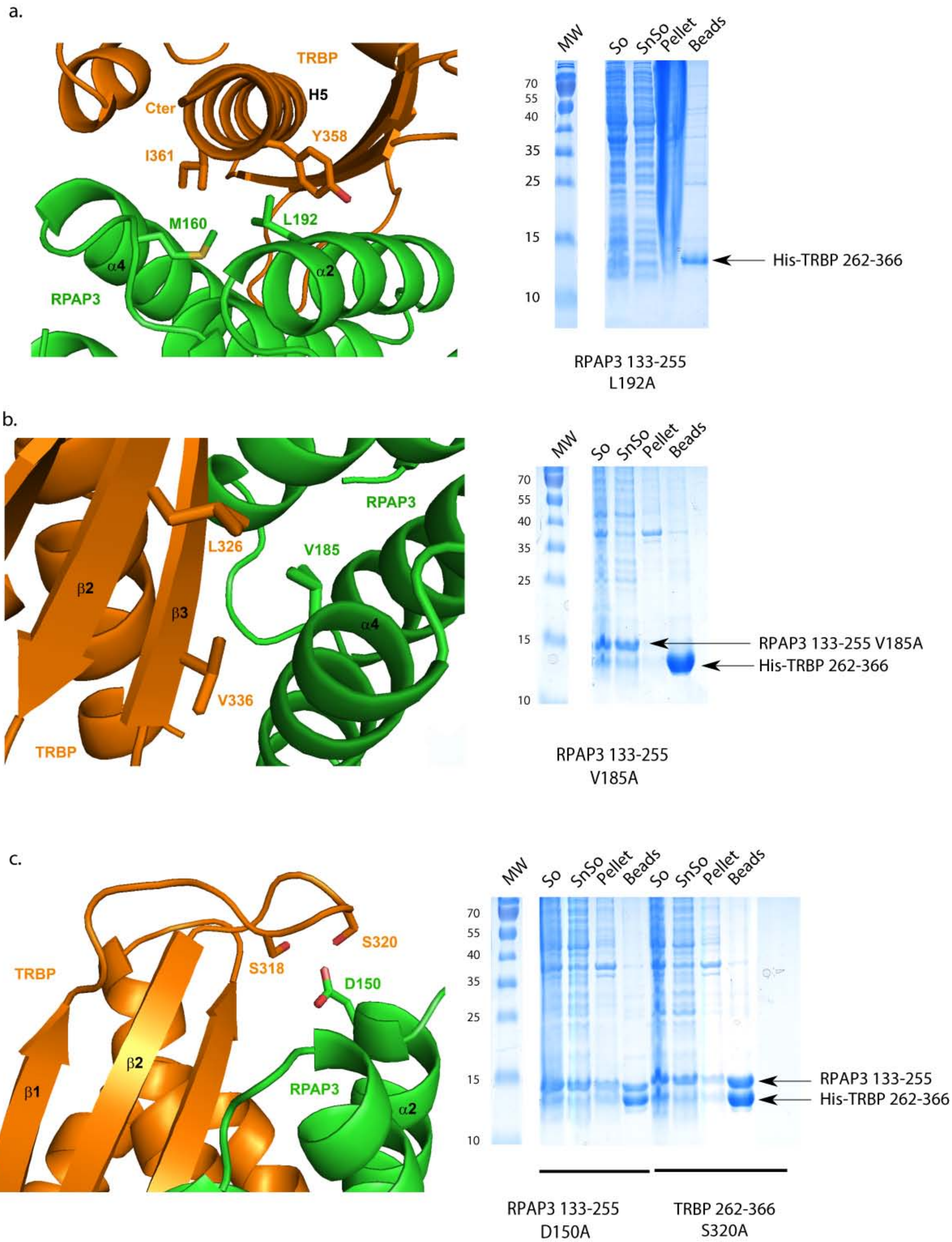
a.
TRBP
H5
Cter
Y358
I361
M160
L192
α4
α2
RPAP3
MW
So
SnSo
Pellet
Beads
70
55
40
35
25
15
10
His-TRBP 262-366
RPAP3 133-255
L192A
b.
RPAP3
L326
V185
β2
β3
α4
V336
TRBP
MW
So
SnSo
Pellet
Beads
70
55
40
35
25
15
10
RPAP3 133-255 V185A
His-TRBP 262-366
RPAP3 133-255
V185A
c.
TRBP
S318
S320
D150
β1
β2
RPAP3
α2
MW
So
SnSo
Pellet
Beads
So
SnSo
Pellet
Beads
70
55
40
35
25
15
10
RPAP3 133-255
His-TRBP 262-366
RPAP3 133-255
D150A
TRBP 262-366
S320A

Supp. Fig. 8

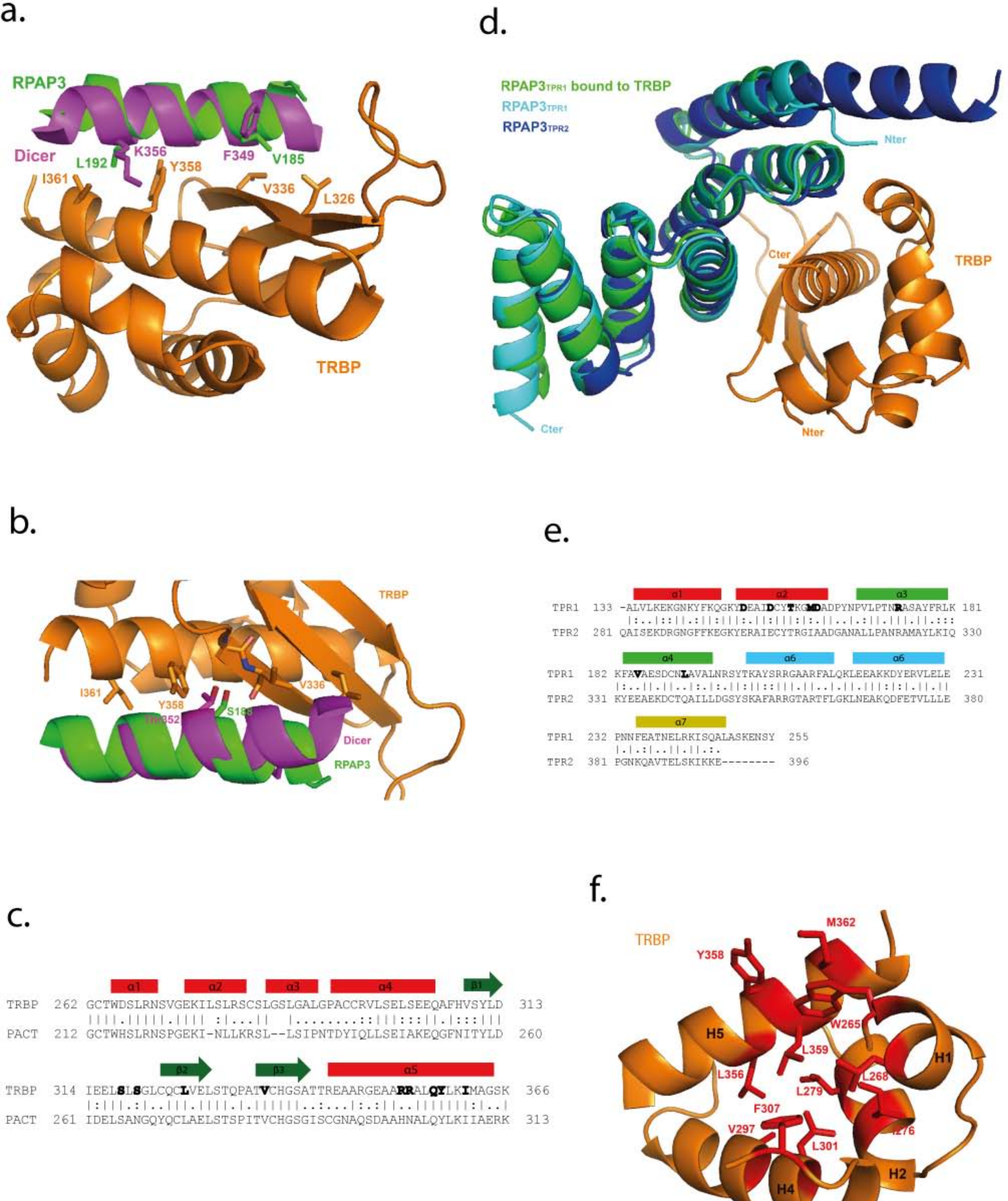
a.
RPAP3
Dicer
L192
K356
Y358
F349
V185
I361
V336
L326
TRBP
d.
RPAP3TPR1 bound to TRBP
RPAP3TPR1
RPAP3TPR2
Nter
Cter
TRBP
b.
TRBP
I361
Y358
S188
V336
Dicer
RPAP3
e.
α1
α2
α3
α4
α6
α6
α7
TPR1 133 -ALVLKEKGNKYFKQGKYDEAIDCYTKGMDADPYNPVLPTNRASAYFRLK 181
TPR2 281 QAISEKDRGNGFFKEGKYERAIECYTRGIAADGANALLPANRAMAYLKIQ 330
TPR1 182 KFAVAESDCNLAVALNRSYTKAYSRRGAARFALQKLEEAKKDYERVLELE 231
TPR2 331 KYEEAEKDCTQAILLDGSYSKAFARRGTARTFLGKLNEAKQDFETVLLLE 380
TPR1 232 PNNFEATNELRKISQALASKENSY 255
TPR2 381 PGNKQAVTELSKIKKE-------- 396
c.
α1
α2
α3
α4
β1
TRBP 262 GCTWDSLRNSVGEKILSLRSCSLGSLGALGPACCRVLSELSEEQAFHVSYLD 313
PACT 212 GCTWHSLRNSPGEKI-NLLKRSL--LSIPNTDYIQLLSEIAKEQGFNITYLD 260
β2
β3
α5
TRBP 314 IEELSLSGLCQCLVELSTQPATVCHGSATTREAARGEAARRALQYLKIMAGSK 366
PACT 261 IDELSANGQYQCLAELSTSPITVCHGSGISCGNAQSDAAHNALQYLKIIAERK 313
f.
TRBP
M362
Y358
W265
H5
L359
H1
L356
L268
L279
F307
V297
L301
H4
H2

Supp. Fig. 9

| | Mutation | position | effect |
|---|---|---|---|
| **RPAP3** | K137A | α1 | Nothing |
| | N141A | α1 | Nothing |
| | D150A | α2 | Nothing |
| | I153A | α2 | Nothing |
| | D154A | α2 | Nothing |
| | T157A | α2 | No expression |
| | D161A | α2 | **No interaction** |
| | D163A | Turn α2-α3 | Nothing |
| | Y165A | Turn α2-α3 | Nothing |
| | N172A | α3 | Nothing – HSP90 Cter binding |
| | R173A | α3 | No expression |
| | Q177A | α3 | Nothing |
| | K181A | Turn α3-α4 | Nothing |
| | K182A | Turn α3-α4 | Nothing |
| | V185A | α4 | **No interaction** |
| | S188A | α4 | Nothing |
| | D189A | α4 | No expression |
| | N191A | α4 | No expression |
| | **L192A** | α4 | No expression |

| | Mutation | position | effect |
|---|---|---|---|
| **TRBP** | S283A | Turn H2-H3 | Nothing |
| | E300A | H4 | Nothing |
| | S310A L312A | β1 | Nothing |
| | I314A | Turn β1-β2 | Nothing |
| | E315A | Turn β1-β2 | Nothing |
| | E316A | Turn β1-β2 | Nothing |
| | S318A | Turn β1-β2 | Nothing |
| | S320A | Turn β1-β2 | Nothing |
| | Q324A | β2 | **No interaction** |
| | L326A | β2 | Nothing |
| | E328A | β2 | **No interaction** |
| | S330A | Turn β2-β3 | Nothing |
| | T331A | Turn β2-β3 | **No interaction** |
| | Q332A | Turn β2-β3 | Nothing |
| | T335A | β3 | Nothing |
| | V336A | β3 | Nothing |
| | H338A | β3 | Nothing |
| | S340A | β3 | Nothing |
| | R354A | H5 | Nothing |
| | R354E | H5 | **No interaction** |
| | Q357A | H5 | **No interaction** |
| | I361A | H5 | Nothing |
| | M362A | H5 | Nothing |

Supplementary Table 1

**Supplementary Figure 1.** Characterization of the TRBP:RPAP3 interaction and absence of PACT:RPAP3 interaction. **a, b.** Yeast two-hybrid control experiments with empty pAS2 or pACT2 plasmids. **c.** Yeast two-hybrid control experiments with the PACT protein expressed from the PACT2 plasmid. **d-f.** Co-expression experiments in *E. coli* between RPAP3 and $His_6$-PACT*,* or subdomains, and between $His_6$-PACT and Dicer subdomains as a control. *Aspecific contaminant band. “So”, “SnSo”, “Beads”, “MW” and “Pellet” relate respectively to the culture sonicate, supernatant sonicate, Talon beads, Molecular Weight markers and sonicate pellet.

**Supplementary Figure 2.** Characterization of the TRBP:RPAP3 interaction. **a.** Controls of co-expression experiments in *E. coli*. LS/HS: low salt, high salt. **b.** $His_6$-TRBP:RPAP3 complexes co-expressed in *E. coli* were eluted from the resin by digestion, and loaded on SDS-PAGE and submitted to gel filtration. See materials and methods for details.

**Supplementary Figure 3.** Characterization of the TRBP:RPAP3 interaction. **a.** *In vitro* GST-pulldown assays using purified recombinant proteins showed that a $His_6$-tagged TRBP copurified with the GST-tagged RPAP3 protein but not with GST alone on Glutathione Sepharose beads. **b.** Co-immunoprecipitation experiments performed in the HEK293T cell line transiently expressing V5-tagged TRBP, with or without RNase A treatment. **c.** Antibody control experiments of the Duolink® assays.

**Supplementary Figure 4.** Characterization of the TRBP:RPAP3 interacting domains. **a.** Yeast two-hybrid control experiments for the RPAP3 subdomains with empty pAS2 or pACT2 plasmids. **b.** Co-expression experiments for the RPAP3 subdomains (TPR1 and TPR2). **c.** Co-expression experiments to determine the narrowed-down domain of TRBP involved in the interaction with the TPR1 domain of RPAP3. Positive interactions are highlighted in red. **d.** Dicer:TRBP co-expression control experiments. “So”, “SnSo”, “Beads”, “MW” and “Pellet” relate respectively to the culture sonicate, supernatant sonicate, Talon beads, Molecular Weight marker and sonicate pellet.

**Supplementary Figure 5.** Structural features of the TRBP:RPAP3 interaction. **a.** Superimposition of the 3D structure of RPAP3 (residues 133-249; in green) from the RPAP3:TRBP complex with the crystal structure of RPAP3 bound to the C-terminal tail peptide (SRMEEVD) of HSP90 (in cyan) (30)

(PDB 4cgv). The 3D structure of RPAP3 in the RPAP3:TRBP complex displays a strictly conserved TPR fold. **b.** Superimposition of the 3D structure of TRBP (residues 263-365; in orange) from the RPAP3:TRBP complex with the crystal structure of TRBP bound to Dicer (in pink) (23) (PDB 4wyq). **c.** Ribbon representation of the 3D structure of TRBP from the crystal structure of the TRBP:RPAP3 complex. This 3D structure of TRBP in the RPAP3:TRBP complex contains a typical dsRBD fold as well as a N-terminal extension.

**Supplementary Figure 6. a.** $^{1}H$-$^{15}N$ HSQC spectrum recorded on $^{13}C/^{15}N$ labeled RPAP3-TPR1:TRBP-dsRBD3 at 950 MHz, 303 K in 10 mM NaPi, pH 6.4, 150 mM NaCl. **b.** Simulated $^{1}H$-$^{15}N$ HSQC spectrum of backbone amide groups of RPAP3-TPR1:TRBP-dsRBD3. Resonances coming from RPAP3-TPR1 are colored in green, resonances coming from TRBP-dsRBD3 are colored in orange. The spectrum was simulated using chemical shift data listed in **(c)**. **d.** TALOS structures derived from NMR data recorded on RPAP3-TPR1:TRBP-dsRBD3 were compared to the secondary structure pattern of the X-ray structure. The two patterns are in good agreement, confirming the similarity between the solution and the crystal states of the complex.

**Supplementary Figure 7.** The RPAP3-TPR1:TRBP-dsRBD3 complex co-elutes with HSPs. **a.** Protein co-expression assays in *E. coli* with the $His_6$-TRBP(262-366):RPAP3(133-255) complex or $His_6$-RPAP3(133-255) and human HSP70 or HSP90-MC. “So”, “SSo”, “Beads” and “Elu” relate respectively to the culture sonicate, supernatant sonicate, Talon beads and elution from the beads with imidazole. The co-purified proteins are indicated with colored arrows. **b.** Protein co-expression assays in *E. coli* with the $His_6$-TRBP:RPAP3 complex or sub-complex and human HSP70 or HSP90-MC. (B) Protein co-expression assays in *E. coli* with $His_6$-RPAP3 (full-length, TPR1 or TPR2) and human HSP70 or HSP90-MC. “So”, “SSo”, “Beads” and “Elu” relate respectively to the culture sonicate, supernatant sonicate, Talon beads and elution from the beads with imidazole. The co-purified proteins are indicated with colored arrows.

**Supplementary Figure 8.** Identification of key residues involved in TRBP:RPAP3 binding interface **a-c.** Hydrogen bonds and hydrophobic residues clusters at the interface of the heterodimer. Co-expressions with each mutant pairs are depicted on the right side of each panel.

**Supplementary Figure 9.** RPAP3 binds to TRBP in the same location as Dicer. **a, b.** The α-helix α4 in RPAP3 and Dicer bind to TRBP in a similar way. Superimposition of the crystal structure of the RPAP3:TRBP complex with the crystal structure of the Dicer:TRBP complex using only atoms from TRBP, with (**a.**) hydrophobic contacts and (**b.**) main-chain (V336) to side-chain (Ser188/Thr352) hydrogen bond at the heterodimer interface. **c.** Sequence alignment comparing TRBP's and PACT's dsRBD3. Residues of TRBP that are involved in the RPAP3:TRBP interface are in bold. **d.** Superimposition of the crystal structure of $RPAP3_{TPR1}$ bound to TRBP (in green and orange) with the crystal structures of $RPAP3_{TPR1}$ (in cyan) and $RPAP3_{TPR2}$ (in blue) bound to the HSP90-tail peptide (PDB 4cgv and 4cgw, respectively) (30). **e.** Sequence alignment of the TPR1 and TPR2 domains of RPAP3. Residues of the TPR1 domain of RPAP3 that are involved in the RPAP3:TRBP interface are bold. **f.** The TRBP structure (residues 263-365) in the TRBP:RPAP3 crystal structure contains a $\alpha/\beta$ sandwich (residues 289-365) typical of a dsRBD fold as well as a N-terminal extension (residues 263-288). Hydrophobic residues that are involved in a large hydrophobic cluster at the interface between the dsRBD and the N-terminal extension of TRBP are displayed.

**Supplementary Table 1.** RPAP3 and TRBP mutations tested.
